# PD-1 endocytosis unleashes the cytolytic potential of check-point blockade in tumor immunity

**DOI:** 10.1101/2024.04.28.591549

**Authors:** Elham Ben Saad, Andres Oroya, Nikhil Ponnoor Anto, Meriem Bachais, Christopher E. Rudd

**Affiliations:** Department of Medicine, University of Montréal, Montréal, QC H3C 3J7, Canada; Centre de Recherche Hopital Maisonneuve-Rosemont (CR-HMR) Montreal, Quebec H1T 2M4; Department of Biochemistry and Molecular Medicine, Montréal, QC H3T 1J4, Canada; Department of Microbiology, Infection and Immunology, Universite de Montreal, Montreal, Quebec, Canada; Division of Experimental Medicine, McGill University, Montreal, QC, H3A 0G4, Canada

**Author notes:** To whom correspondence should be addressed-Christopher E. Rudd. Equal shared first author (Elham Ben Saad, Andres Oroya).

**Keywords:** PD-1, checkpoint blockade, endocytosis, tumor immunotherapy

## Abstract

PD-1 immune checkpoint blockade (ICB) is now a promising first-line treatment for many cancers. While the steric blockade of PD-1 binding to its ligand plays a role, the role of internalisation in promoting the efficacy of ICB has not been explored. In this study, we show that PD-1 internalisation also contributes by unlocking the full cytolytic potential of ICB in cancer immunotherapy. We found that anti-mouse and human PD-1 downregulate a subset of PD-1 surface receptors on T-cells with high-density surface PD-1 leaving T-cells with intermediate expression resistant to further internalisation. Down regulation was seen on both CD4 and CD8 cells but was maximally effective on CD8 effector cells. In human T-cells, nivolumab outperformed pembrolizumab in terms of rate and efficacy. We also found that PD-1 internalisation depended on bivalent antibody (Ab)-induced crosslinking, while monovalent Ab sterically blocked PD-1 without inducing endocytosis. Immunologically, while both monovalent and bivalent Ab limited B16-PD-L1 tumor growth, bivalent Ab was significantly more effective. In molecular terms, while both antibodies increased granzyme B (GZMB) expression in CD8+ cytolytic T-cells, the induction of the second key cytolytic pore-forming mediator, perforin, was dependent on the blockade and internalisation mediated by bilavent anti-PD-1. Our findings unveil a novel mechanism in checkpoint blockade where steric blockade combined with the removal of PD-1 from the cell surface by endocytosis can complement and optimize therapy. The targeting of PD-1 internalisation holds promise for enhancing anti-tumor immunity and improving PD-1 checkpoint blockade therapy.

**Graphical Abstract:** 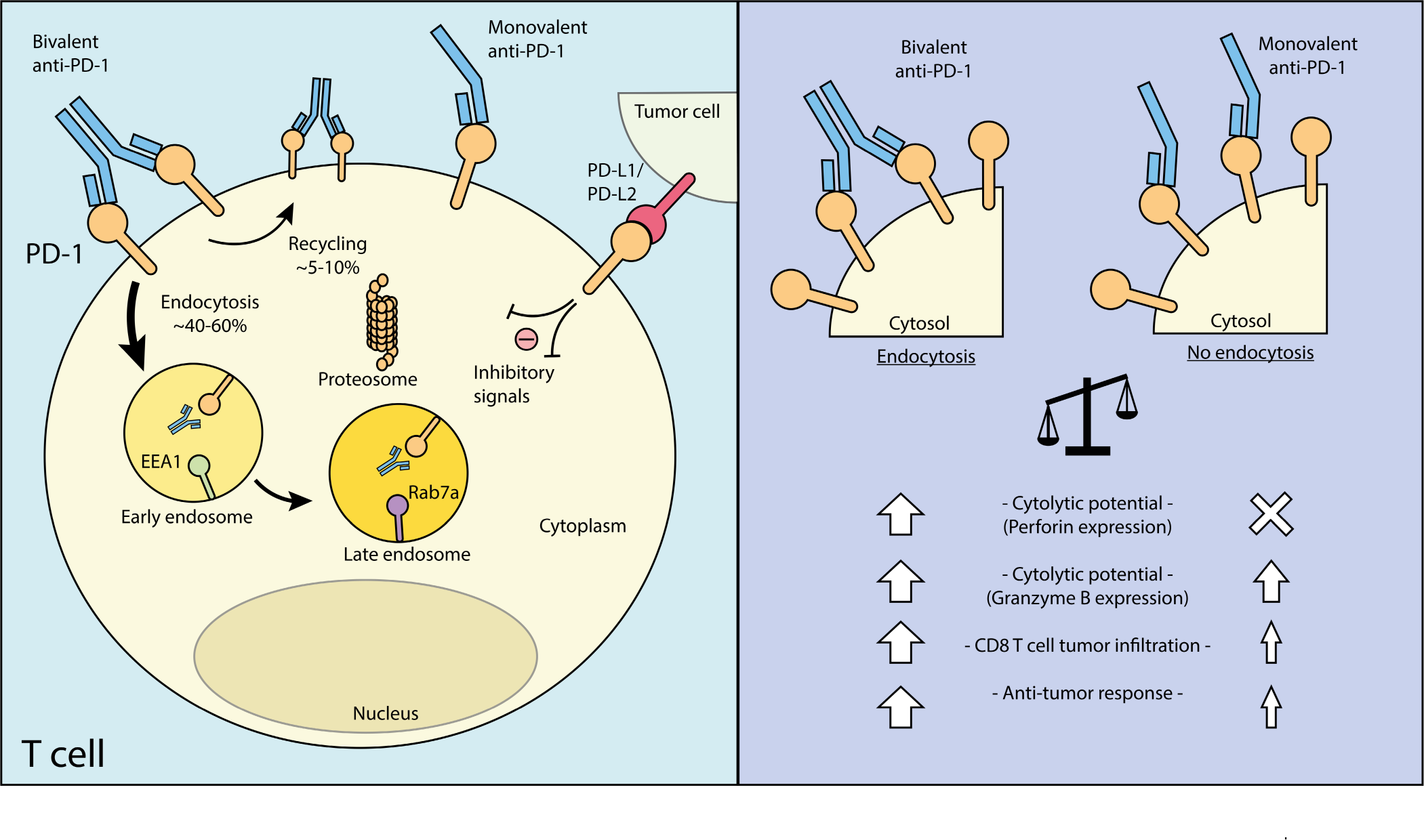

**In brief:** Ben Saad et al define the mechanism of PD-1 inhibitory endocytosis and show that the removal of surface PD-1 by endocytosis plays a role in complementing and optimizing checkpoint blockade. Targeting PD-1 internalisation holds promise for enhancing anti-tumor immunity and improving the efficacy of PD-1 checkpoint blockade therapy.

## Introduction

The programmed cell death 1 (PD-1; PDCD1) coreceptor, belonging to the B7 gene family, plays a role in the inhibition of T-cell function ^1,2^. The co-receptor binds to the programmed cell death ligands 1 and 2 (PD-L1/L2) which are expressed on both lymphoid and nonlymphoid cells^3,4^. Following T-cell activation, PD-1 is expressed and contributes to T-cell exhaustion^5–7^. Anti-PD1, in turn, can reverse exhaustion and restore T cell functionality ^8,9^. In this context, the blockade of PD-1 and PD-L1 with monoclonal antibodies in immune checkpoint blockade (ICB) has shown therapeutic benefits in patients with various cancers. These tumors include non-small cell lung carcinoma (NSCLC), melanoma, and bladder cancer^10–13,11^. Response rates have been observed with anti-PD-1 alone, or in combination therapy with anti-cytotoxic T-lymphocyte–associated antigen 4 (CTLA-4) ^11,14–16^. The two most commonly used antibodies, Nivolumab and Pembrolizumab are effective despite having distinct binding affinities to different epitopes on PD-1 ^17,18^. PD-1 expression on tumor-infiltrating CD8^+^ T cells correlates with impaired effector cell function ^19^, while PD-L1 expression on tumors can facilitate escape from immunity ^1^ and serves as a prognostic factor^20^. Nevertheless, despite the success of ICB, a significant proportion of patients are not cured, emphasizing the ongoing need for a better understanding of checkpoint blockade mechanisms to facilitate the development of more effective therapies.

Over the years, there has been a focus on understanding the intracellular signaling pathways the govern PD-1 function in T cells. We and others showed that T cells are activated by tyrosine phosphorylation activation cascade involving the protein tyrosine kinases p56^lck^ and ZAP-70^21,22^. p56^lck^ binds to coreceptors CD4 and CD8 ^23,24^ and phosphorylates immune receptor activation motifs (ITAM) needed for ZAP-70 recruitment to the TcR–CD3 complex ^25^. In this context, the co-ligation of PD-1 can dephosphorylate substrates of TCR and CD28 induced signaling^26,27^. The cytoplasmic domains of PD-1 have immunoreceptor tyrosine-based inhibition motifs (ITIMs) and immunoreceptor tyrosine-based switch motifs (ITSMs)^28^. These domains serve as docking sites for SH2 domain-containing protein tyrosine phosphatase (Shp)-2 (encoded by the Ptpn11 gene) and its homologous counterpart, Shp-1 ^29^. Both phosphatases contain SH2 domains that interact with phospho-tyrosine motifs on receptors ^30–32^. Further, mutation of the YESM motif disrupts the negative signaling mediated by PD-1 in B and T-cells ^3,33–41^. However, surprisingly, PD-1 is still functional in shp-2 deficient mice ^42^. Moreover, CD8+ T cells lacking both phosphatases have been reported to still differentiate into exhausted cells and to respond to PD-1 blockade ^43^. On the other hand, Pdcd1 expression is regulated by a diverse interplay of transcription factors, including Nuclear Factor of Activated T cells (NFAT), Forkhead Box Protein O1 (FoxO1), Notch, Activator Protein 1 (AP1), and B-lymphocyte Maturation Protein 1^21,44–47^. More proximally, we showed that glycogen synthase kinase 3 (GSK-3) can downreglate PD-1 expression via the upregulation of the transcription factor, Tbet which inhibits PD-1 and LAG-3 transcription^48–50^. GSK-3 is a serine/threonine kinase that is active in resting T cells and becomes inactivated with T-cell activation^51^. Further, we showed that the inhibition of GSK-3 with small molecule inhibitors (SMIs) can suppress tumor growth, comparable to PD-1 antibody treatment ^48,49^. Recent studies have shown that PD-1 is also expressed on myeloid cells where it may also contribute to the effects of ICB in the control of tumor growth^52,53^.

The ligation of receptors by ligand or antibody causes their endocytosis and removal from the surface of cells. Receptors differ in their ability to undergo endocytosis ^54^. In immune cells, endocytosis begins with kinase-mediated signaling, followed by the translocation of the microtubule-organizing center (MTOC) and endosomes to the immunological synapse ^55^. Receptors have been reported to use clathrin-coated pits or caveolae for internalisation ^56,57^, enter intracellar endocomes and may undergo proteolytic degradation in lysosomes and/or be recycled to the cell surface ^58^. Receptor endocytosis is also influenced by its association with intracellular proteins. For instance, the binding of CD4 to the protein-tyrosine kinase p56^lck^ can inhibit clathrin-dependent entry (38). Further, we showed that CD28 binding to phosphatidyl inositol 3 kinase (PI-3K) affects the efficiency of CD28 internalisation^59^. Further, CTLA-4 is released to the cell surface by orphan adaptors TRIM and LAX from the trans-Golgi network ^60,61^. The movement of vesicles containing the LAT protein is associated with protein micro-cluster organization ^62^, while the intra-flagellar transport system protein IFT20 facilitates LAT and TCR transport back to the cell surface ^63^. In the case of PD-1, steady-state endocytosis is mediated by the E3 ligase FBXO38 ^64^.

Surprisingly, despite the importance of checkpoint blockade in sterically preventing PD-1 binding to PDL-1, the role of PD-1 internalisation in promoting the efficacy of ICB has not been fully explored. In this paper, we define aspects of PD-1 internalisation and show for the first time that the removal of surface PD-1 by endocytosis complements and optimizes checkpoint blockade. We show that the internalisation of PD-1 requires receptor crosslinking which leads to a more potent cytotolytic program than seen with mere steric blockade of PD-1. Targeting PD-1 internalisation holds promise for enhancing anti-tumor immunity and improving the efficacy of PD-1 checkpoint blockade therapy.

## Results

### Anti-PD-1 binding induces the endocytosis of PD-1 on T-cells

To investigate PD-1 internalisation, we initially conducted an internalisation analysis on human T-cells, following a methodology previously utilized for CD28 ^59^. Human T-cells were activated with anti-CD3/CD28 for 2 days to induce PD-1 expression, rested for 24 hours before commencing the down-modulation studies. In the first approach, cells were incubated with anti-PD-1 (clone J110) (5ug/ml) at 4°C on ice for 30 minutes, washed to remove unbound antibody, and then incubated at 37°C for varying times. Subsequent staining of surface anti-PD-1 with AF488-anti human IgG enabled the detection of remaining surface-bound anti-PD-1 molecules (Fig. 1A). This revealed a reduction of 15-20% in surface PD-1 levels by 15 minutes, 20-25% within 30 minutes, and 35-40% within 60 and 90 minutes (Fig. 1B). Notably, a significant portion of PD-1 was resistant to down-modulation, aligning with the concept of spare receptors or density-dependent endocytosis ^65^. The down-modulation of anti-CD28, as previously described ^59^, demonstrated similar levels of internalisation for both CD28 and PD-1 in response to their respective antibodies (Fig. 1C).

**Figure 1:**
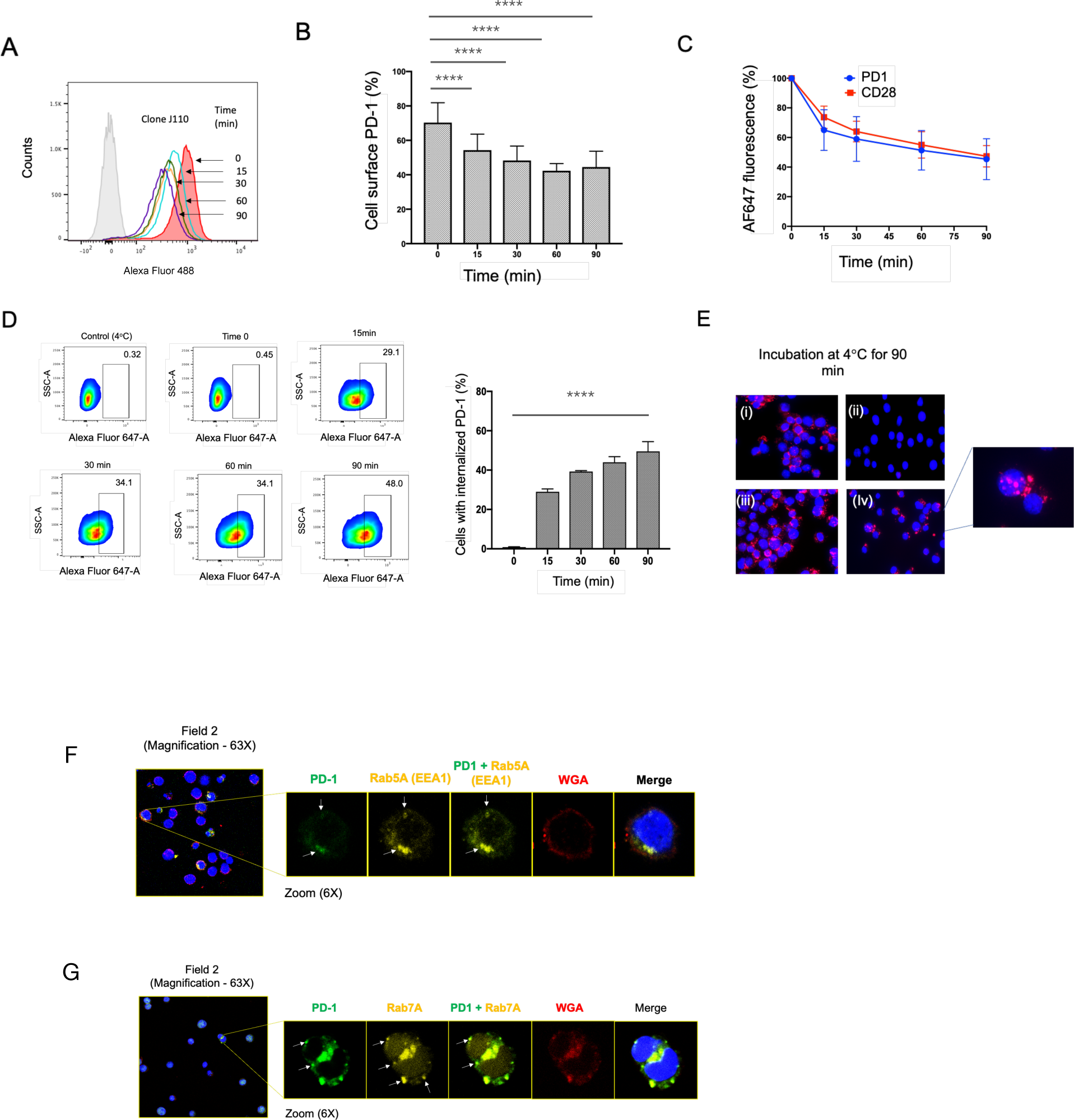
Internalisation of PD-1 Induced by Anti-PD1 on Human T-Cells. **Panel A:** Standard FACs profile demonstrating that anti-human PD1 (J110) induces the internalization of a portion of PD1 over a time period of 0-90 minutes. Human T-cells were activated with anti-CD3/CD28 for 2 days to induce PD-1 expression, followed by a 24-hour rest period before conducting internalization studies. The cells were pre-incubated with saturating amounts of anti-PD-1 (clone J110) at 4°C, washed, and then incubated at 37°C for varying durations. Subsequently, samples were stained with a secondary antibody (AF647 anti-mouse IgG or AF488-anti-human IgG) to detect the presence of surface-bound anti-PD-1 (n=4). **Panel B:** Time course analysis of PD1 internalization presented as a percentage of the total original staining. The J110 antibody induced the down-regulation of 40% of surface PD1 within 30 minutes of incubation (n=4). **Panel C:** Time course comparison of PD1 and CD28 internalization from the surface of human T-cells (n=3). **Panel D:** Time course showing internalization using an alternate technique involving an acidified dissociation solution. Human T-cells were incubated with PE-conjugated anti-PD-1 J110 for varying durations at 37°C and then treated with an acidified dissociation solution to remove surface-bound anti-PD-1. Left panel: Anti-PD1 induced the internalization of 45-50% of the receptor within 30 minutes, followed by 50-60% within 70-90 minutes (n=4). Right panel: FACs panels indicating acquired resistance to dissociation. **Panel E:** Immunofluorescence microscopy demonstrating the internalization of PD1 from the surface of T-cells. While antibody bound to cells at 4°C was easily removed with acid treatment (panels iii vs. i), an immunofluorescence signal was observed in cells subjected to incubation at 37°C followed by acid treatment (n=3). **Panel F**: Internalised anti-PD1 antibody complexes co-localize with the early endosome marker EEA1 (Rab5a). C57BL6J mice-derived splenic T-cells were incubated with anti-PD-1 (clone RMP1-14) incubated briefly with acidic PBS (pH-2) to remove cell surface antibody followed by fixation and the staining with either anti-EEA1. Left panel: overview of multiple cells; right panels: PD-1, EEA1, PD1-EEA1 overlap, WGA to stain cell surface; merge including DAPI staining for the nucleus. **Panel G:** Internalised anti-PD1 antibody complexes co-localized with the late endosome marker Rab7a. As above.

Using a second alternate approach, human T-cells were incubated with the PE-conjugated anti-PD-1 J110 at 37°C for different durations. Surface anti-PD-1 was removed using an acidified dissociation solution, and the presence PE-conjugated antibody was measured to assess internalisation (Fig. 1D). The low residual staining for T-cells at 4°C following acid treatment showed the effectness of the acidified solution in removing surface-bound PD-1 (right panel). Employing this method, we it was observed that 29% of PD-1 internalisation by 15 minutes, increasing to 34% by 30-60 minutes and up to 48% by 90 minutes (left FACs plots and right histogram). The internalisation of antibody-PD-1 complexes was confirmed by confocal microscopy (Fig. 1E). Notably when cells that had been incubated with anti-PD-1 at 4°C for 60min followed by treatment with acid showed no internalisation (panel ii vs. i). By contrast, cells that had been incubated at 37°C for 60 minutes showed a significant amount of internalised signal (i.e., iv vs. iii). The down-modulation was seen in CD4 and CD8 T-cells (Fig. S1). These findings demonstrate that antibody binding to PD-1 triggers its internalisation from the surface of human T-cells.

Further, we assessed whether the internalised anti-PD1 antibody complexes co-localized with the early endosome marker EEA1 (Rab5a) or the late endosome marker Rab7a (Fig. 1E, F). C57BL6J mice-derived splenic T-cells were incubated with anti-PD-1 (clone RMP1-14) for 1h followed by treatment with acidic PBS (pH-2) to remove cell surface antibody followed by fixation and the staining with either anti-EEA1 or Rab7a Abs. In the case of EEA1, many cells showed internalised PD-1 in large vesicles that co-localized with EEA1 (Fig. 1E). More than 75% of cells with internalised PD-1 co-localized with anti-EEA1 (see Table S1). Further, internalised anti-PD1 also colocalized with the late endosome marker Rab7a (Fig. 1F). In this case, we documented that 64% of cells with internalised PD-1 showed co-localized Rab7A (table S1). These data suggested that anti-PD-1 complexes are internalised via the endosomal pathway.

### Nivolumab is more effective in inducing PD-1 internalisation than pembrolizumab

We were next interested in the behaviour of nivolumab and pembrolizumab since the two therapeutic reagents are used in cancer immunotherapy ^11,15^. Previous studies have shown that nivolumab and pembrolizumab target distinct epitopes on the PD-1 molecule ^66–68^. To confirm this, we conducted an experiment where pembrolizumab (1μg/ml) was initially bound to T-cells followed by anti-human Alexa Fluor-488 at 4°C, washing and the addition of nivolumab at fivefold higher levels (5 μg/ml) for 30 minutes. Notably, the addition of nivolumab did not displace the binding of pembrolizumab, thereby supporting the notion that they bind to distinct sites on the PD-1 molecule (Fig. 2A). Further, we found that nivolumab exhibited higher binding levels when compared to pembrolizumab (Fig. 2B). This difference was evident in the higher mean fluorescence intensity (MFI) for nivolumab (MFI of 754) in contrast to pembrolizumab (MFI of 487) (upper and lower panels, respectively). Nevertheless. despite targeting different epitopes on the PD-1 molecule, both antibodies were able to induce internalisation of PD-1 from the surface of T-cells within 30-120 minutes (Fig. 2C, upper and lower FACs profiles). As seen in the lower histogram, Nivolumab was generally more effective, causing a more significant reduction within 30 minutes. Further, this was also observed over a range of antibody concentrations from 10ug/ml to 1ug/ml (Fig. 2D). Further, we found the rate of endocytosis of nivolumab was more rapid than pembrolizumab when measured over 0-30 minutes (Fig. 2E). Further, the combination of nivolumab and pembrolizumab did not increase in efficacy of PD-1 down-regulation relative to nivolumab alone (Fig. 2F). These findings indicated that nivolumab was generally more effective in the downregulation of PD-1 than pembrolizumab.

**Figure 2:**
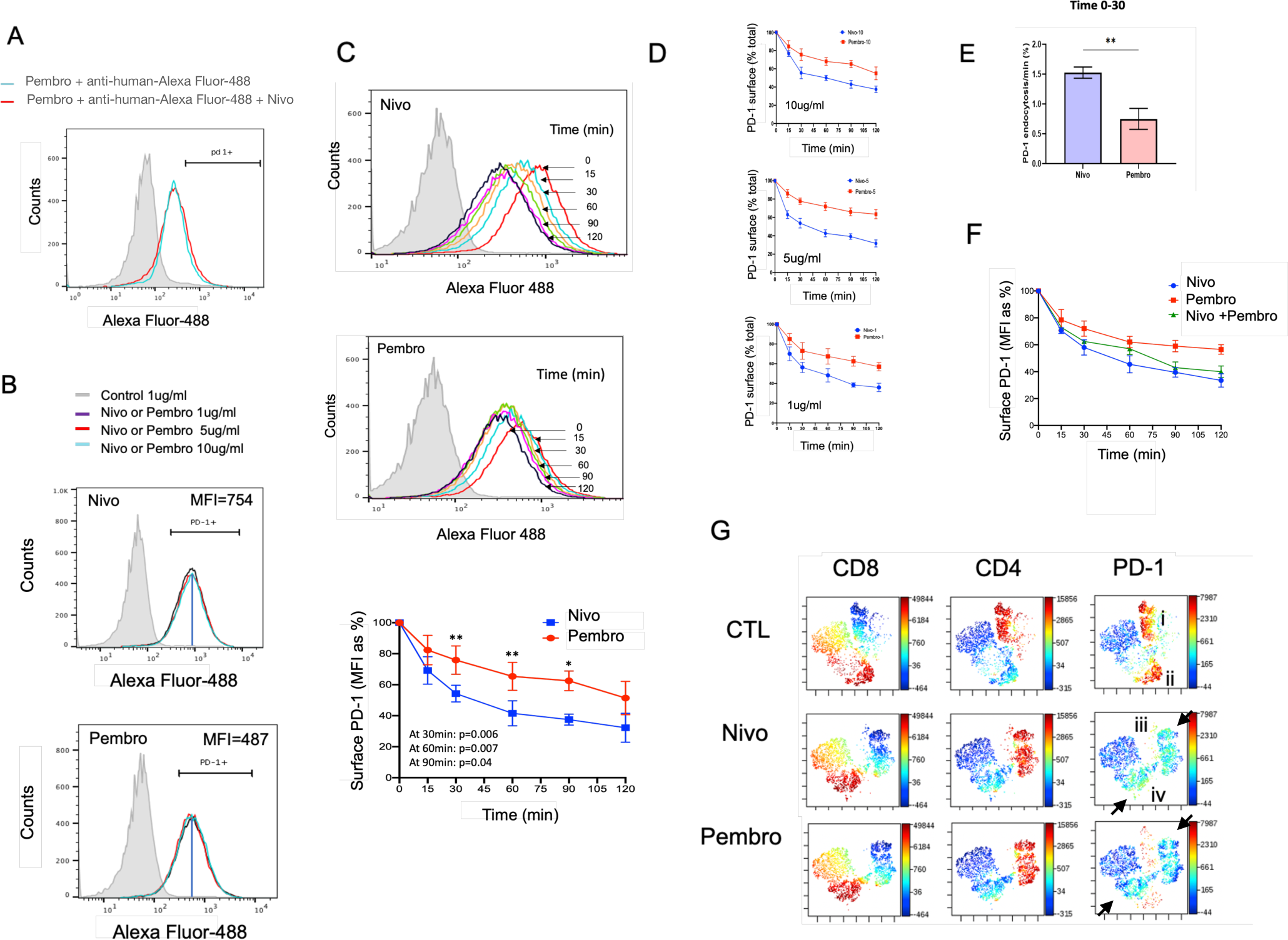
Nivolumab is more effective than Pembrolizumab causing the conversion of high-density PD-1 to low-density T-cells. **Panel A**: Confirmation of distinct binding sites on PD-1 for Nivolumab and Pembrolizumab. The binding of Pembrolizumab (Alexa Fluor-488) (1ug/ml) is not displaced by unlabeled Nivolumab (1ug/ml). (n-=2) **Panel B:** Nivolumab binds more to T-cells than Pembrolizumab. Nivolumab (1-10ug/ml) showed a higher mean fluorescence intensity (MFI) on T-cells compared to Pembrolizumab. Nivolumab has an MFI value of 754, while Pembrolizumab has an MFI of 487 (n=3). Panel C: Both Nivolumab and Pembrolizumab induce PD-1 internalization. T-cells are activated with anti-CD3/CD28 to express PD-1, followed by a rest period. Cells are then exposed to saturating amounts of Nivolumab or Pembrolizumab at 4oC, washed to remove excess antibody, and incubated at 37oC for various times. Staining with secondary antibodies detects surface-bound anti-PD-1. Upper and middle panels show the time-dependent internalization of PD-1 induced by Nivolumab and Pembrolizumab. The lower panel displays a histogram of the time course of PD-1 down-regulation by both antibodies (n=4). **Panel D:** Histograms indicating the loss of Nivolumab or Pembrolizumab-induced internalization over time (0-120min) at different antibody concentrations (1, 5, 10ug/ml) as a percentage of the original binding (MFI) (n=3). **Panel E:** Histogram representing the rate of endocytosis as a percentage relative to total binding over 6-minutes (n=2). **Panel F:** Nivolumab induces PD-1 internalization more rapidly than Pembrolizumab, including when used in combination (n=4) **Panel G:** viSNE profiles show the loss of PD-1 high-intensity clusters. Control (CTL) samples show the presence of a high-intensity PD-1 cluster on CD4 and CD8+ clusters (upper panel, clusters i and ii, respectively). Nivolumab (middle panel) and pembrolizumab (lower panel) incubation resulted in the loss of both clusters accompanied by the appearance of two new clusters with intermediate PD-1 expression (clusters iii and iv). These data show that anti-PD1-induced endocytosis causes the loss of high-density PD-1 from the surface of T-cells.

We next employed multi-dimensional ViSNE analysis to assess potential effects on subsets of T-cells (Fig. 2G). Distinct clusters of CD4 and CD8 T-cells were evident (see red). Futher, anti-PD-1 identified a subset of CD4 and CD8 cells in clusters i and ii, respectively. Incubation with nivolumab for 90-minute at 37°C resulted an marked reduction in staining in both CD4 and CD8 cell clusters, leaving new clusters of cells with low levels of staining. This was seen by the light blue staining in CD4 cluster iii and CD8 cluster iv (middle panels; see arrows). Pembrolizumab also caused this shift, albeit less completely than nivolumab, as evidenced by the presence of some remaining cells with high PD-1 expression in clusters i and ii. These findings show visually that the anti-PD-1 antibodies convert T-cells with high PD-1 surface expression to cells with low-intermediate PD-1 expression.

### Anti-PD-1 internalizes PD-1 on effector memory and effector subsets

Our results confirmed that both nivolumab and pembrolizumab decreased PD-1 expression in CD4+ T-cells (Fig. 3A, upper and lower panels, respectively) and CD8+ T-cells (Fig. 3B, upper and lower panels, respectively) (also see Fig. S1). CD4 cells routinely showed higher levels of expression than CD8+ T-cells and in both cases, nivolumab bound at higher levels than pembrolizumab. This was also indicated by the MFI values for CD4+ T-cells (MFI=790 for nivolumab and 520 for pembrolizumab) and CD8+ T-cells (MFI=348 for nivolumab and 230 for pembrolizumab (Fig. 3C). Down-modulation by both antibodies left a lower residual population of PD-1 on human CD8 than CD4+ T-cells.

**Figure 3:**
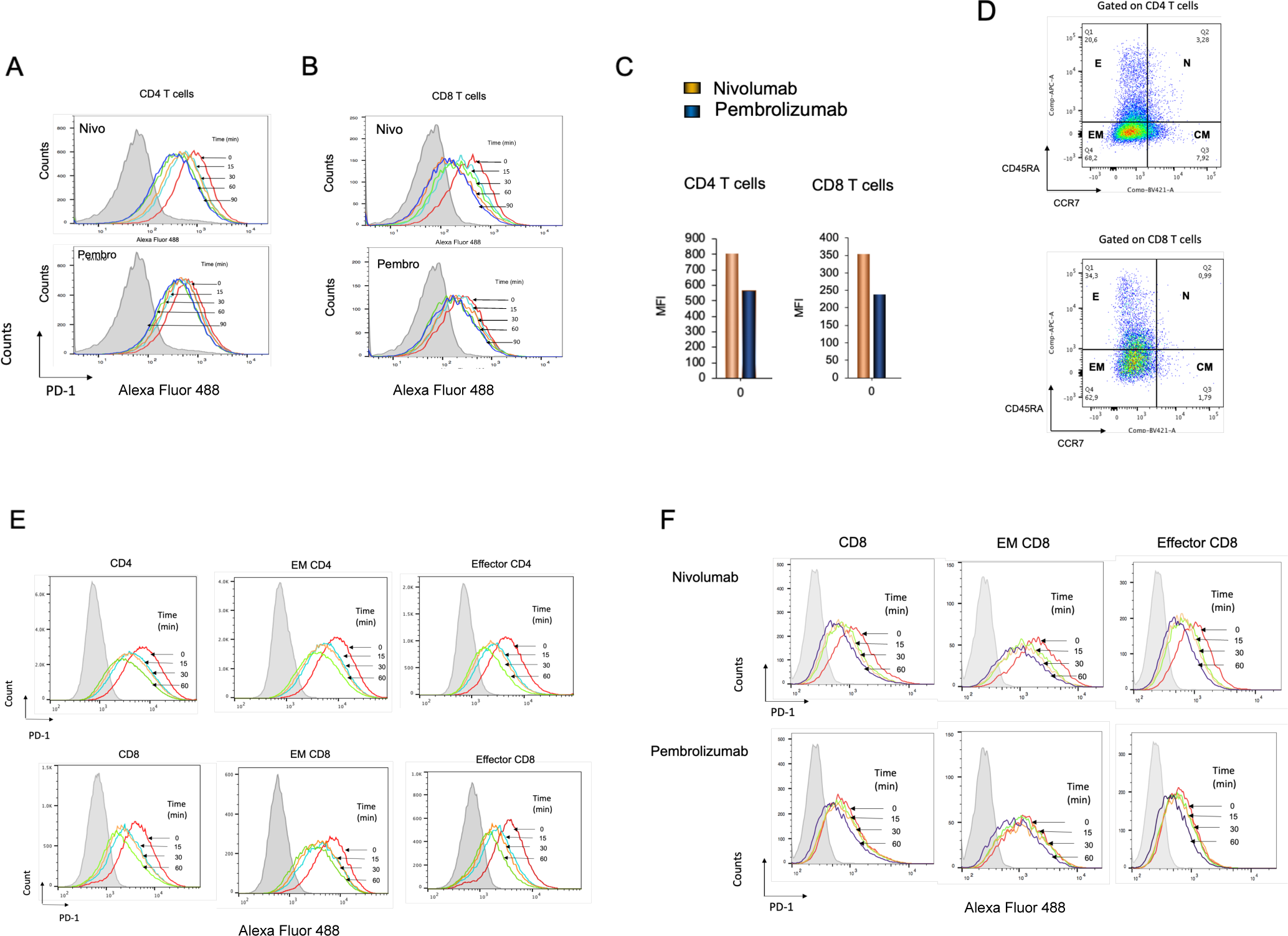
Anti-PD1 downregulates PD-1 from different T-cell subsets. **Panel A:** Histogram profiles showing the down-regulation of PD-1 expression on CD4+ T-cells by nivolumab (upper panel) and pembrolizumab (lower panel) (n=4). **Panel B:** Histograms demonstrating the down-regulation of PD-1 expression on CD8+ T-cells by nivolumab (upper panel) and pembrolizumab (lower panel) (n=4). **Panel C:** Nivolumab binds at higher levels to activated human CD4 and CD8 T-cells than pembrolizumab, as demonstrated by the histogram depicting mean fluorescence intensity (MFI) values. The left panel illustrates the MFI values for nivolumab and pembrolizumab binding to human CD4 T-cells, while the right panel shows the MFI values for the same antibodies binding to CD8+ T-cells (n=3). **Panel D:** Gating strategy for effector memory and effector CD4 and CD8+ human T-cells. The figure displays FACs profiles employed to identify effector memory CD4 and CD8+ T-cells, based on the staining of CD45RA and CCR7 markers. The experiment was conducted with a sample size of four (n=4). Upper panels: CD4 cells; Lower panels: CD8 T-cells **Panel E:** FACs analysis of PD-1 expression on effector memory (EM) and effector (E) CD4 and CD8+ T-cells. The figure illustrates the impact of nivolumab on the down-regulation of PD-1 expression in effector memory and effector CD4 (upper panels) and CD8+ (lower panels) T-cells (n=4). **Panel F:** FACs analysis of nivolumab and pembrolizumab induced downregulation of PD-1 on effector memory (EM) and effector (E) CD8+ T-cells over time. Upper panel: Nivolumabcinduced PD-1 down-regulation; Lower panel: Pembrolizumab induced PD-1 down-regulation.

Different T-cell subsets were evident based on gating strategies with anti-CD45RA and CCR7 staining (Fig. 3D). Due to a limited number of cells available, we used nivolumab to assess effects on effector memory (EM) and effector CD4+ and CD8+ T-cells (Fig. 3E). CD4+ cells showed higher levels of expression on EM and effector CD4 cells than on the same subsets of CD8+ T-cells. To our surprise, EM showed higher levels of staining than effector T-cells. Nivolumab exhibited a greater affinity for binding to EM cells compared to effector T-cells. Consequently, this downregulation led to a smaller residual surface staining of PD-1 on effector T-cells. With a focus on CD8+ T-cells, a similar effect was observed in comparing the binding and downregulation of PD-1 by nivolumab and pembrolizumab (Fig. 3F). Both antibodies bound at higher levels to effector memory and than effector cells. Further, the down-regulation of PD-1 resulted in the smallest level of residual PD-1 expression on effector CD8+ T-cells. These findings are consistent with previous studies that attribute the anti-tumoral effects of anti-PD-1 to the effector CD8 subset ^69,70^.

### Antibody-induced PD-1 endocytosis is clathrin/caveolar-independent

Various cell surface internalisation pathways like clathrin-mediated endocytosis, caveolar endocytosis, and micropinocytosis deliver their cargoes to intracellular endosomes^54,71^. To investigate this in the context of PD-1, T-cells were initially co-incubated at 37°C with hypotonic sucrose, which inhibits clathrin-mediated endocytosis and other caveolar mediated pathways. The hypotonic solution disrupts the osmotic gradients and water flux required for the invagination process^71^. As in Figure 1D, T-cells were incubated with the PE-conjugated anti-PD-1 J110 at 37°C followed by antibody removal using an acidified dissociation solution, and the remaining PE-conjugated antibody was measured to assess internalisation. Cells incubated with sucrose showed lower levels of internalized PD-1. While 48% of T-cells internalised PE-tagged anti-PD-1 by 30min, only 32% of cells internalsed the antibody in the presence of sucrose (Fig. 4A). This pattern was similar to the reduction in anti-CD28 from 42% to 32%, as we previously reported (Fig. 4B) (38).

**Figure 4:**
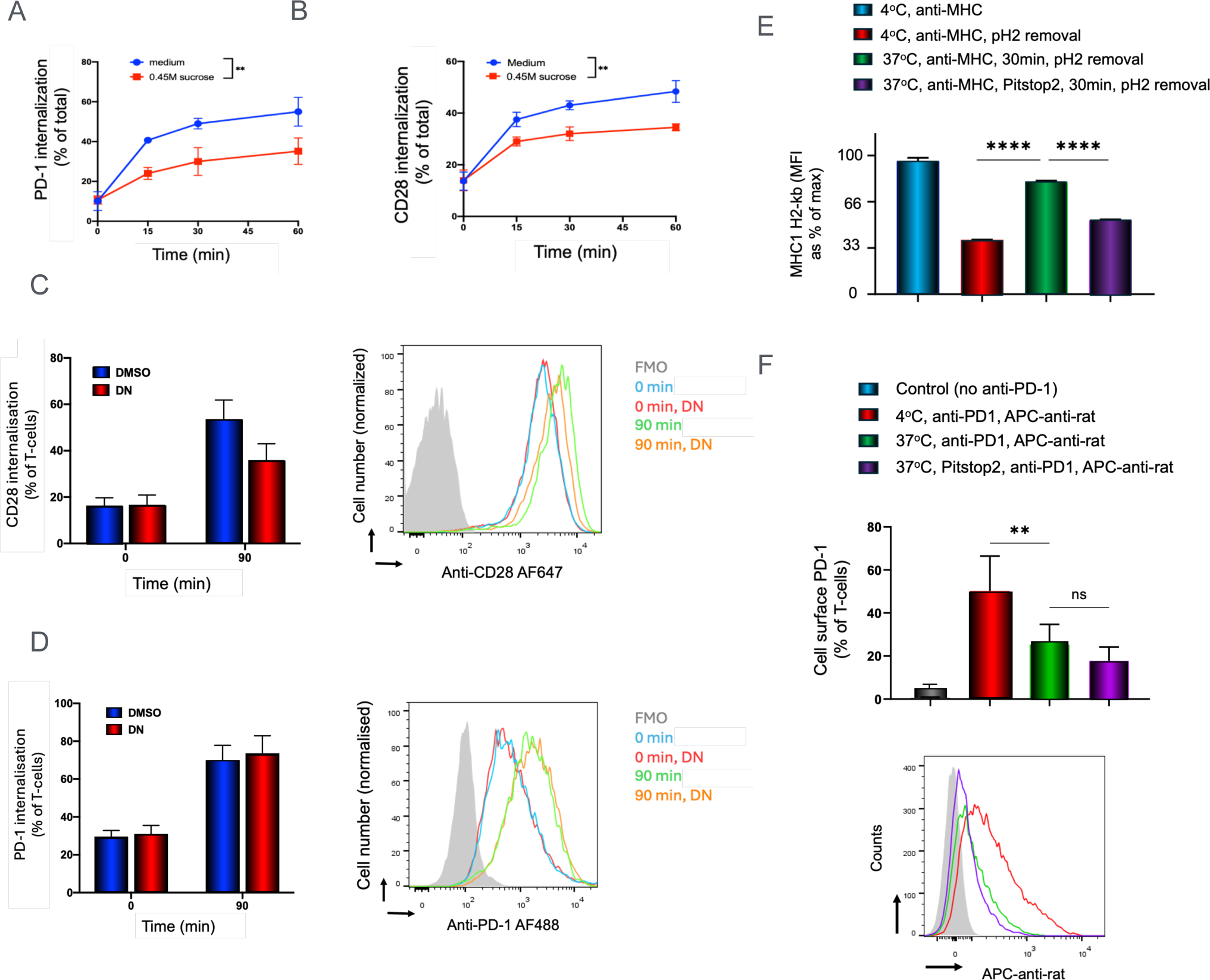
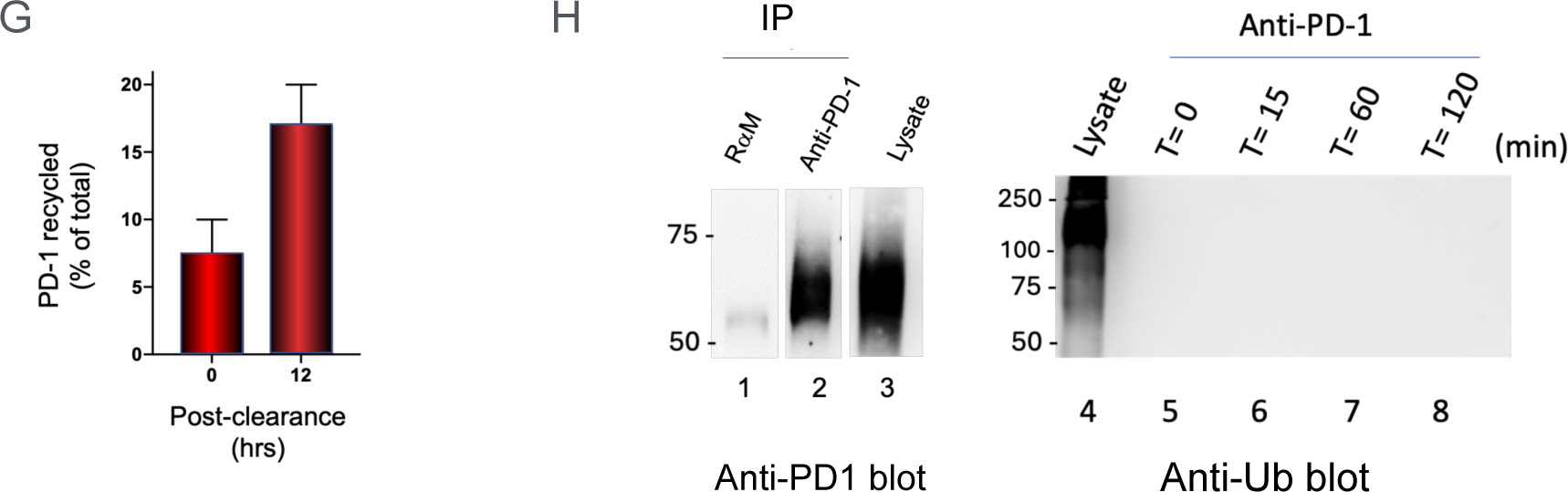
Anti-PD1 induced internalisation and recycling is proteasome-dependent but independent of clathrin, caveolae, and ubiquitination. **Panel A:** PD-1 induced internalisation is inhibited by hypotonic sucrose (n=3). In this assays, as outlined in Figure 1D, Human T-cells were activated with anti-CD3/CD28 for 2 days to induce PD-1 expression. Following washing away stimulatory antibodies and a rest period of 24 hours, T-cells were incubated with the PE-conjugated anti-PD-1 J110 at 37°C for different durations. Surface anti-PD-1 was then removed using an acidified dissociation solution, and the remaining PE-conjugated antibody was measured to assess internalisation. Cells incubated with sucrose showed lower levels of internalized PD-1. **Panel B:** CD28 induced internalisation is inhibited by hypotonic sucrose (n=3). The internalization of CD28 was measured using the same approach as in Panel A. Cells incubated with sucrose showed lower levels of internalized CD28. **Panel C:** Anti-CD28 induced internalisation is inhibited by incubation with Dynsore (n=3). CD28 internalisation was assessed in activated human T-cells for 90min in the absence or presence of DN following the protocol as in panel B. Left panel: histogram showing the percent PD-1 internalization; right panel: FACs profile. **Panel D:** Anti-PD1 induced internalisation was not affected by incubation with Dynsore (DN)(n=3). PD-1 internalisation was assessed in activated human T-cells for 90min in the absence or presence of DN as in panel A.. Left panel: histogram showing the percent PD-1 internalization; right panel: FACs profile. **Panel E:** Pitstop inhibits MHC class 1 antigen induced internalization (n=2). As outlined in Figure 1A-C, mouse activated T-cells were then incubated with anti-PD-1 (clone RMP1-14) at 4°C on ice for 30 minutes, washed to remove unbound antibody, and then incubated at 37°C for varying times. **Panel F:** Pitstop fails to affect anti-PD1 induced internalization (n=2). As outlined in Figure 1A-C, mouse activated T-cells were then incubated with anti-PD-1 (clone RMP1-14) at 4°C on ice for 30 minutes, washed to remove unbound antibody, and then incubated at 37°C for varying times. Subsequent staining of surface-bound anti-PD-1 with AF488-anti rat enabled the detection of remaining surface-bound anti-PD-1. Left panel: FACs profile; right panel: histogram showing the percent PD-1 internalization. **Panel G:** Internalized anti-PD1-PD1 complexes can be recycled to the surface of T-cells. Activated human T-cells were incubated with the anti-PD-1 J110 at 37°C for 60min followed by surface anti-PD-1 removal using an acidified dissociation solution. Cell were then left to incubate in RPM1 1640 media with 10% FCS and antibiotics overnight followed by staining of cells with PE-conjugated anti-mouse antibody to measure surface PD-1. **Panel H:** Anti-PD1 internalisation occurs without detectable ubiquitination (n=3). Anti-PD1 internalisation occurs without detectable ubiquitination (n=3). Activated human T-cells were bound by anti-PD-1 at 4°C followed by an incubation from 0 to 120 min at 37°C. Cells were then solubilized in Triton X-100 containing lysis buffer, followed by the addition of anti-PD-1 or IgG and precipitation using Protein A Sepharose beads. Rabbit anti-mouse and anti-PD1 precipitates as well as cell lystaes were subjected to immunoblotting for PD-1 (lanes 1-3). Similarly, cell lysates (lanes 4) or anti-PD-1 precipitates over times of 0 to 120min were subjected to anti-Ubiquitin (Ub) blotting (lanes 4-8).

Given this, we also conducted endocytosis assays using cells co-incubated with the dynamin inhibitor dynasore (DN) (Fig. 4C, D). DN inhibits clathrin-mediated endocytosis at the transition from a half and fully formed pit to an endocytic vesicle ^72,73^ but also been reported to inhibit the caveolar pathway^73^. Anti-CD28-AF647 or anti-PD-1 AF488 was incubated with human T-cells for 90min followed by the measurement internalised antibody-antigen complexes. Control (i.e., DMSO) treated cells showed an increase in antibody upake in 56% of T-cells (histogram, left panel) and FACs plot (right panel). The incubation with dynasore partially inhibited anti-CD28 internalisation, showing uptake in only 38% of T-cells (Fig. 4C). By contrast, DN had no detectable effect on the internalization of PD-1 when assessed at 90 minutes (Fig. 4D) and other time points (data not shown). The internalisation of PD-1 was seen in 78% of T-cells when assayed at 90min following incubation at 37°C (histogram: left panel) and FACs plot (right panel). The incubation with DN had no effect on the level of internalised PD-1. These data showed that the incubation with a specific inhibitor of clathrin and caveolar mediated internalisation did not affect the endocytosis of PD-1 in human T-cells.

We also tested the effect of a second inhibitor Pitstop2 on internalisation (Fig. 4E). Pistop2 is an inhibitor of clathrin-dependent and independent endocytosis^74^ As a control, activated murine T-cells were assessed for the internalisation of MHC class 1 antigen, a surface antigen known to depend on clathrin. T-cells were incubated with the PE-conjugated anti-murine MHC class 1 at 37°C for 30min in the presence and absence of 15uM Pitstop2 followed by removal using an acidified dissociation solution, and the remaining conjugated antibody was measured to assess internalisation. MFI values were expressed as a % of the maximal binding. From this, after a 4°C binding on ice, acidified PBS reduced the flourescent intensity of staining to 26% of T-cells. A 30min incubation at 37°C resulted in an increase in the MFI level of internalised anti-MHC1 increased to 72%. As expected, an incubation with Pitstop2 inhibited the iincrease in internalised anti-MHC1. These data confirmed that Pitstop2 could inhibit MHC class 1 internalisation.

Given this, we next assessed the effects of Pitstop2 on anti-PD-1 internalisation (Fig. 4F). In this case, due to the absence of an effective tagged anti-mouse PD-1 antibody, we followed the internalisation protocol in Fig. 1-A-C. Anti-PD-1 (RMP1-14) was preincubated at 4°C on ice for 30 minutes with T-cells, washed to remove unbound antibody, and then incubated at 37°C for 30min. Subsequent staining of surface-bound anti-PD-1 with AF488-anti rat which recognises the rat RMP1-14 for detection of the remaining surface-bound anti-PD-1. As expected, anti-PD1 incubation at 37°C reduced the presence of surface from 51 to 22% in the absence of inhibitor. In the presence of Pitstop2, anti-PD1 induced a similar level of internalisation (i.e. 19%; histogram (upper panel) and FACs plot (lower panel). We also observed no difference in the MFI of expression. Given that neither Dynasore nor Pitstop2 affected PD-1 internalisation, using human or mouse cells with different assays, our data indicate that anti-PD-1 induced internalisation not being mediated by a conventional clathrin or caveolae-mediated endocytosis pathway. In this context, a new mobile endocytic network connecting clathrin-independent receptor endocytosis to recycling has been reported which promotes T cell activation ^75^. In either case, PD-1 endocytosis was seen to be eventually incorporated into early and late endosomes as shown by EEA1 (Rab5a) and Rab7a colocalization (Fig. 1E, F).

To evaluate the potential re-expression of anti-PD-1-PD-1 complexes on the cell surface following internalization (Fig. 4F), the cells were treated with the antibody for 90 minutes to allow for internalization. Cell surface antibody was removed through acid treatment, and the cells were then incubated overnight for 12 hours. To identify the presence of recycled anti-PD-1 on the cell surface, the cells were re-stained with anti-IgG. The results revealed that 16% of the initial anti-PD-1 signal was detected as complexes. As a control, anti-IgG was used to detect complexes after acid treatment, which exhibited only 6.5% staining compared to the original non-acid-treated cells. This result indicated that some 9-10% of anti-PD1 complexes could be recycled to the surface of T-cells.

Finally, we investigated whether anti-PD-1-induced endocytosis involved PD-1 ubiquitination (Fig. 4G). FBXO38, an E3 ligase has been reported to mediate Lys48-linked poly-ubiquitination of PD-1^64^. Activated human T-cells were subjected to immunoprecipitation with either control rabbit anti-mouse (lane 1) or anti-PD-1 (lane 2) followed by blotting with anti-PD-1. Cell lysate extracts were also run and subjected to anti-PD1 blotting (lane 3). Anti-PD-1 blotting identified a clear band at 45-50Kda in anti-PD-1 precipitates and cell lysates corresponding to PD-1. In the same vien, we also used an anti-ubiquitin antibody to blot against cells lysates (lane 4), and anti-PD1 precipitates at time 0 (lane 5) and over a time course of anti-PD-1 down-regulation (lanes 5-8). Anti-ubiquitin showed the presence of numerous bands across a spectrum of Mrs as expected ^76^. From this, it was evident that anti-Ub antibody could recognize an spectrum of uniquitinated proteins in the cell lysate (lane 4). Despite this, the immunoprecipitation of PD-1 from cells incubated with anti-PD1 over a time course of 0 to 120min failed to show the presence of bands identified by anti-Ub blotting (lanes 5-8). (for Full length blots Fig. S2). These findings suggest that PD-1 ubiquitination is unlikely to be the primary mechanism underlying anti-PD-1-induced internalization in pre-activated T-cells.

### PD-1 crosslinking is needed for receptor internalization

Since current therapeutic antibodies are generally bivalent, it was of interest to determine whether receptor dimerization was needed to induce PD-1 internalisation. In this context, we generated a monovalent form of anti-mouse and human antibody composed Fabc′ chains according to the manufacturer’s instructions (Pierce Chemicals, Rockford, IL) and as we previously published for anti-CD28 ^59^. Monovalent and bivalent anti-mouse J43 showed similar binding to preactivated mouse T-cells expressing PD-1 (Fig. 5A, left panel). viSNE plots confirmed binding CD4 (middle panel) and CD8+ T-cells (right panel).

**Figure 5:**
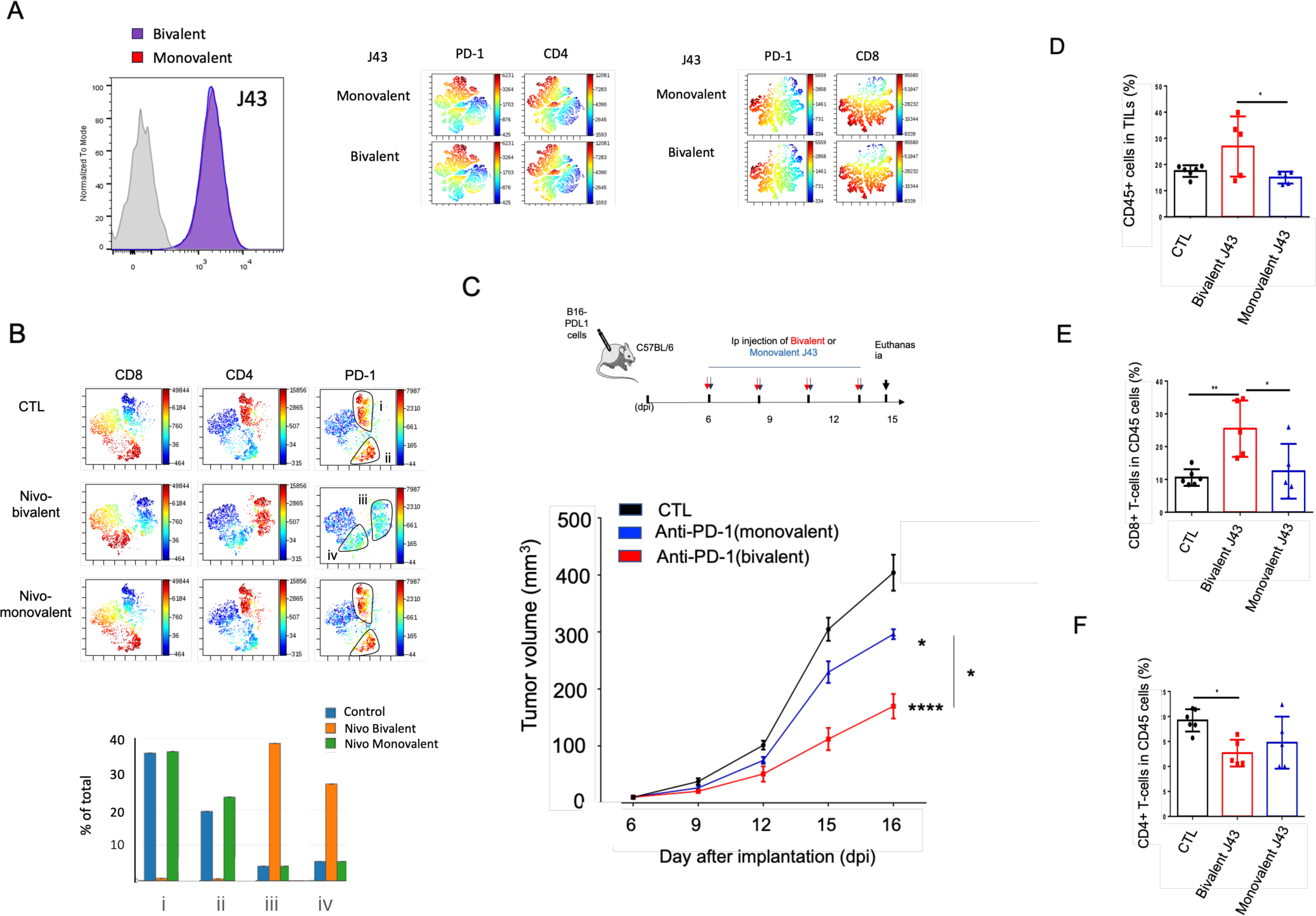
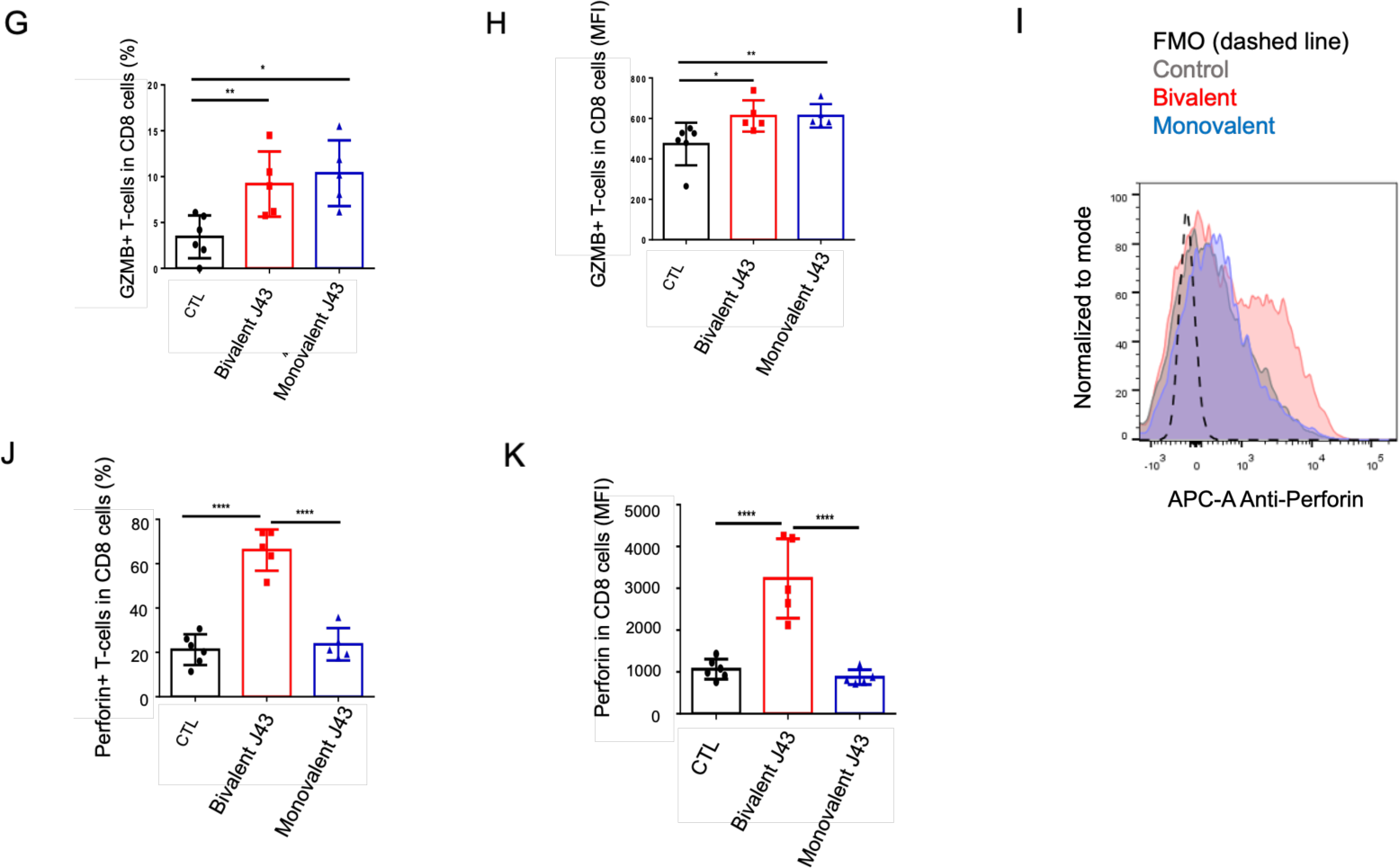
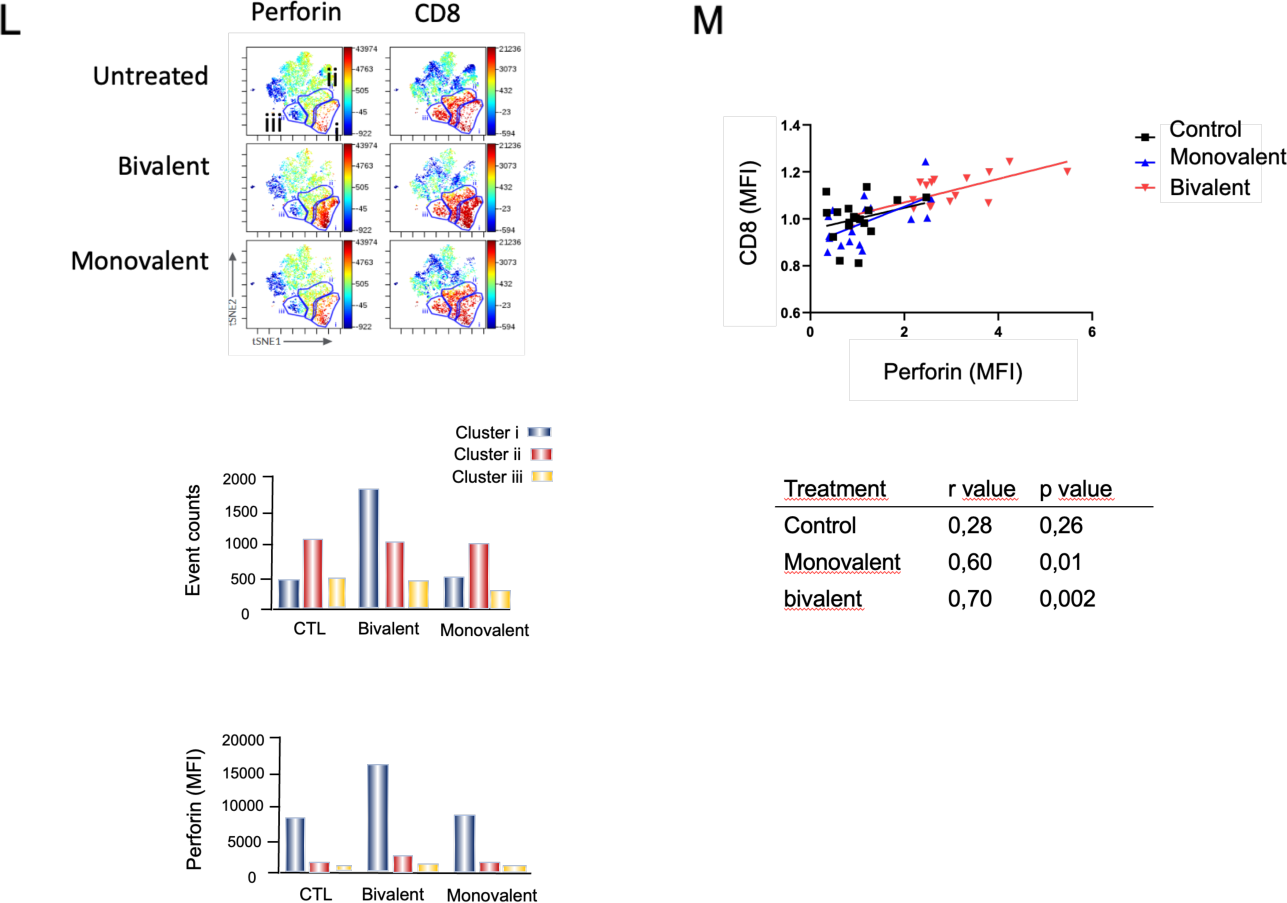
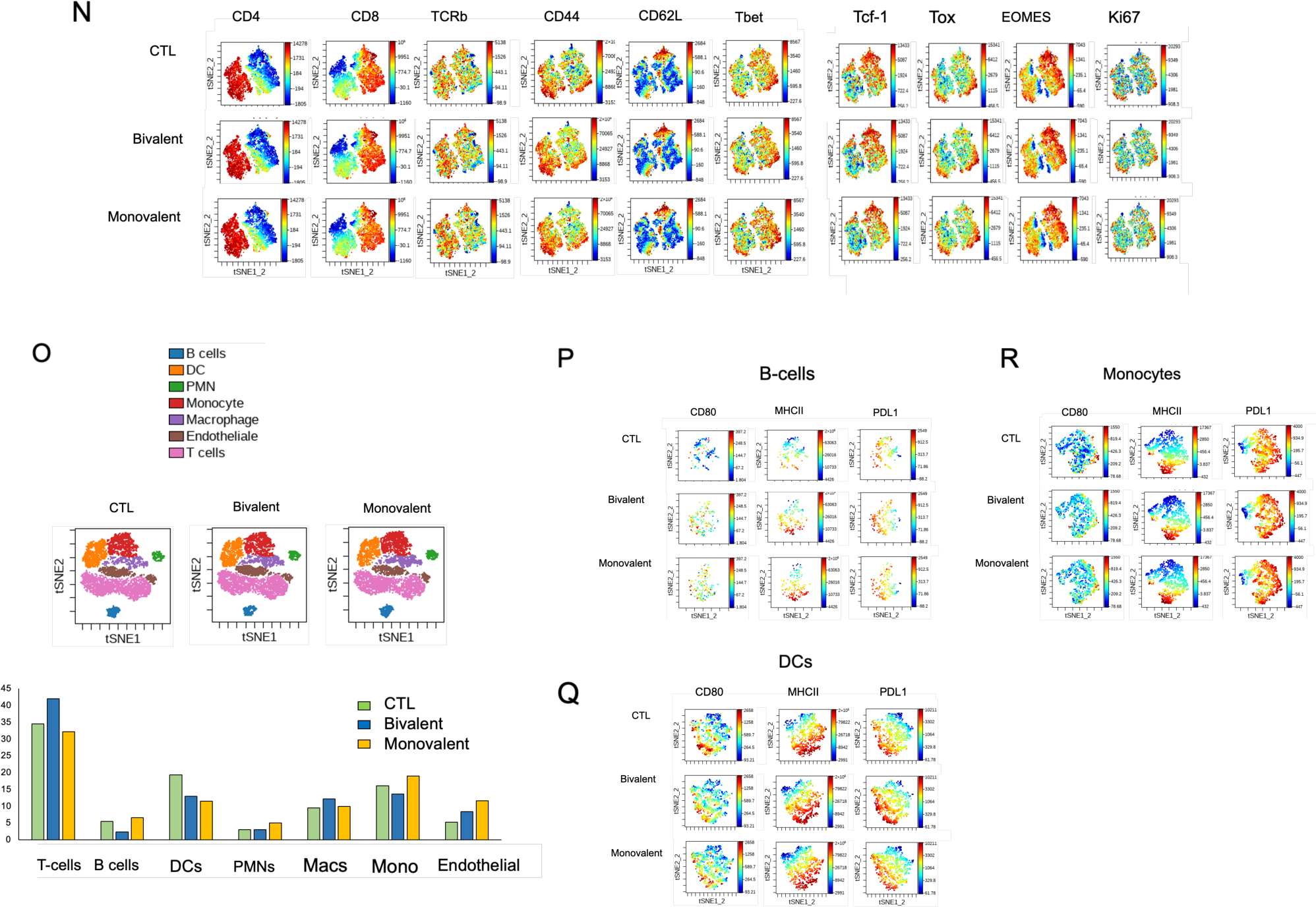
The relative effects of monovalent vs bivalent antibody on tumor elimination. Bivalent anti-PD-1 preferentially affects the expression of perforin for CD8+ cytolytic function. **Panel A:** Monovalent and bivalent J43 bind to CD4 (upper panel) and CD8 (lower panel) T-cells. 1ug/ml of monovalent and bivalent was incubated for 5min (blue) and 30min (dark red) with cells before washing and analysis by flow cytometry (n=2). **Panel B:** viSNE profiles showing that bivalent but not monovalent nivolumab down-regulates PD-1 on human activated human T-cells. Upper panel: Bivalent nivolumab control staining of T-cells for 30min at 4°C. The staining identified two clusters of high PD-1 (clusters i and ii) on CD4 and CD8 cells respectively (control). Middle panel: Bivalent nivolumab incubation with T-cells for 60min at 37°C. viSNE profile shows that clusters i and ii were largely reduced and replaced by the presence of new clusters with low-intermediate PD-1 expression (clusters iii and iv). Lower panel: monovalent nivolumab was incubated with T-cells for 60 minutes at 37oC and showed no downregulation of anti-PD-1 staining (n=5). **Panel C**: Histogram showing the growth of B16-F10 tumors in mice in the absence and presence of biovalent and monovalent anti-PD-1. Upper panel: Regime for the treatment of mice with B16-PD-L1+ tumors; Lower panel: While monovalent and bivalent J43 anti-PD-1 limited B-16 F10 tumor growth, bivalent anti-PD1 had a greater effect (n=4). **Panel D:** Bivalent but not monovalent J43 promotes the presence of increased numbers of CD45+ T-cells in tumors (n=3). **Panel E:** Bivalent but not monovalent J43 therapy increases the presence of CD8+ T-cells within the CD45+ TIL population (n=3). Panel F: Bivalent but not monovalent J43 therapy reduces the relative presence of CD4+ T-cells within the CD45+ TIL population. **Panel G:** Both bivalent and monovalent J43 therapy promote an increased proportion of GZMB expressing cells within the CD8+ TIL population. **Panel H:** Both bivalent and monovalent J43 therapy promote an increased expression of GZMB in the CD8+ TIL population. **Panel I:** FACs profile of the effects of bivalent but monovalent J43 therapy on the expression of perforin within the TIL population. **Panel J:** Bivalent but not monovalent J43 therapy promotes a major increase in the percentage of CD8+ T-cells expressing perforin within the CD8+ TIL population. **Panel K:** Bivalent but not monovalent J43 therapy promotes a major increase in the level of expression (MFI) of perforin within the CD8+ TIL population. **Panel L:** viSNE profiles showing the effects of bivalent J43 antibody crosslinking and endocytosis on therapy on perforin expression on CD8+ TILs. Upper panel: viSNE profiles showing the presence of 3 clusters (i-iii) in which bivalent J43 increases the signal in cluster i relative to the untreated and monovalent therapy. Middle panel: histogram shows the increase in the event count (number of cells) in cluster 1 caused by bivalent J43; lower panel: histogram showing the increase in the MFI for perforin expression in cluster 1 caused by bivalent J43. **Panel M:** Regression profile showing the correlation between CD8 and perforin expression induced by both monovalent and bivalent anti-J43. Lower table: The r value and p values for bivalent J43 were stronger than for monovalent J43. **Panel N:** viSNE profiles showing that bivalent J43 did not affect other aspects of TILs as defined by a comparison of CD44, CD62L, Tbet, TCF-1, Tox. Eomes and Ki67 expression. **Panel O:** Bivalent J43 does not have major effects on other TIL cells. Upper tSNE profile showing the representation of different populations of TILs in response to bivalent vs. monovalent engagement of PD-1. Bivalent showed an increase in numbers of T-cells with a reduction in B-cells and little effect on the presence of PMNs, macrophages (Macs), monocytes (Mono), and endothelial cells. **Panel P**: viSNE profiles showing that bivalent and monovalent therapy both increased the presence of MHC class II and PDL1 expression on B-cells.c **Panel Q:** viSNE profiles showing that bivalent and monovalent therapy showed a trend in decreasing CD80 expression while having little effect on MHC class II or PDL1 expression on DCs. **Panel R:** viSNE profiles showing that bivalent and monovalent therapy showed bivalent and monovalent therapy had little effect on the monocytic populations.

**Figure 5:**
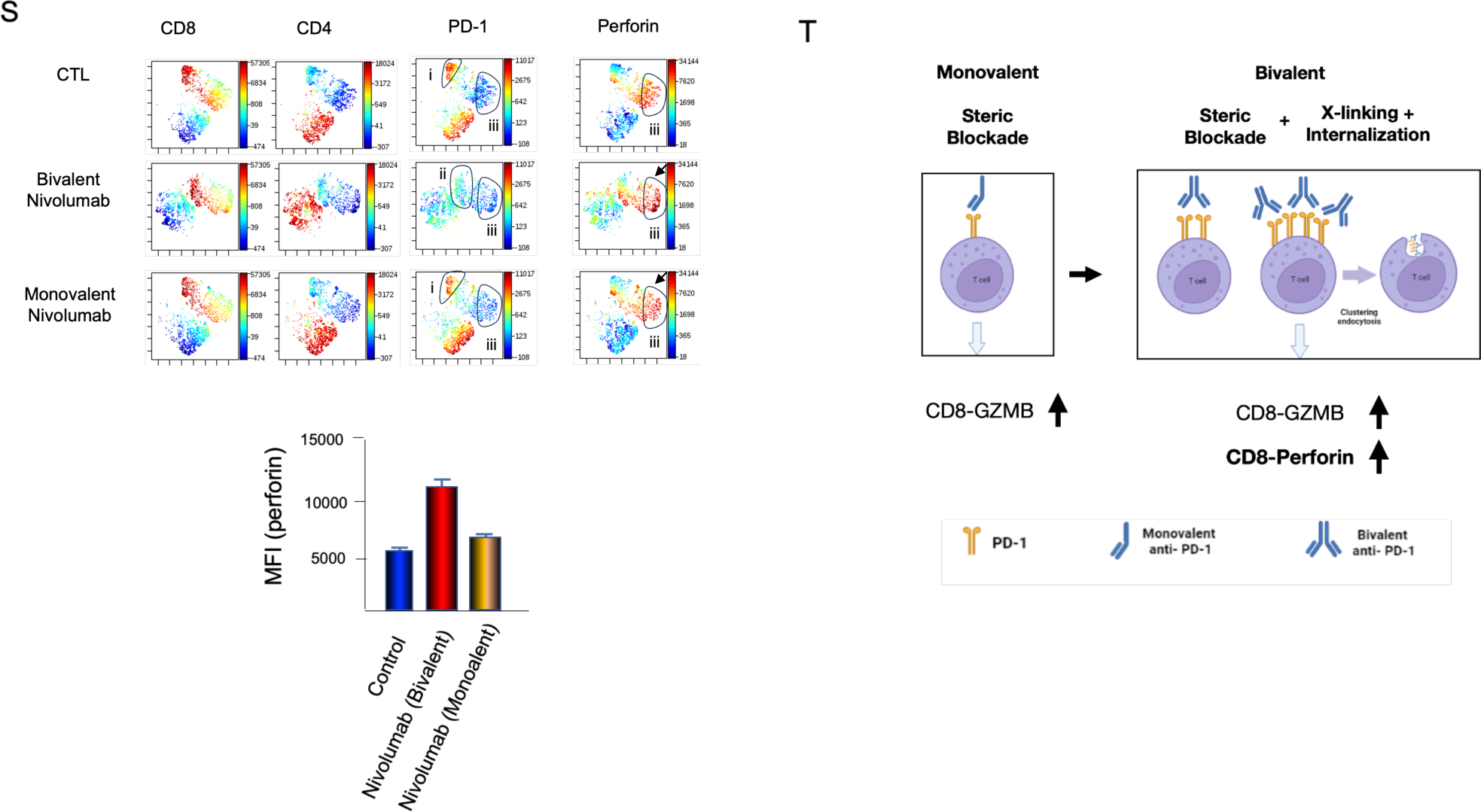
Perforin expression in human CD8+ T-cells requires bivalent Nivolumab. **Panel S:** Human T-cells were initially activated for 24 hours, followed by a period of rest and incubation with either monovalent or bivalent Nivolumab. Afterward, the expression of perforin was analyzed. viSNE profiles revealed the presence of two distinct clusters (i and iii) of CD8+ T-cells. As seen in untreated control cells, cluster i expressed high levels of PD1 and no perforin. Cluster iii expresses high levels of perforin and low PD1. The downregulation of PD-1 with bivalent antibodies resulted in the loss of cluster I, the emergence of a new cluster ii with intermediate PD-1 levels, and an increase in the signal intensity for perforin in cluster iii. This change was not seen with monovalent Nivolumab which instead resembled the pattern in control samples. Lower panel: a histogram illustrating the increase in mean fluorescence intensity (MFI) of perforin in response to bivalent antibody treatment. **Panel T**: **Model illustrating the differential effects of steric blockade on CD8+ T-cells.** The model compared the impact of monovalent steric blockade (achieved through anti-PD1 antibodies) with steric blockade combined with internalization (achieved through bivalent anti-PD1 antibodies). The model shows that steric blockade with either monovalent or bivalent anti-PD-1 can regress tumors and induce an increase in GZMB expression. By contrast, bivalent induced anti-PD-1 internalisation can regress tumors more effectively and an increase in the expression of perforin. This increase in perforin was observed in both *in vivo* and *in vitro* assays, showing a requirement for steric blockade combined with receptor crosslinking and internalisation. These data show that anti-PD-1 induced endocytosis unleashes the cytolytic potential of check-point blockade in tumor immunity. It further suggests that the targeting PD-1 endocytosis can optimize the potential of checkpoint blockade for anti-tumor immunity.

In addition, we showed that bivalent but not monovalent anti-PD-1 induced internalisation. As an example, when incubated at 37°C for 30 min, bivalent nivolumab identified two distinct clusters (cluster i and ii) on CD8 and CD4 cells respectively (upper panel). However, when incubated at 37°C for 60min bivalent antibody caused the disappearance of clusters i and ii leading to the appearance of two new clusters (iii and iv) with intermediate-low PD-1. By contrast, the monovalent antibody showed a pattern that was similar to that seen for bivalent control staining at 4°C (also shown in the histogram, lower panel). These findings are consistent with the notion that the downregulation of PD-1 is dependent on the dimerization and crosslinking of PD-1 by antibodies.

Given this, we next examined the relative effects of monovalent versus bivalent antibodies on tumor regression (Fig. 5C). PDL1+ B16 F10 melanoma cells were implanted in mice and allowed to develop for 6 days before receiving injections of monovalent or bivalent anti-PD-1 (J43) every 2 days (upper and lower panels). Both antibodies were administered at saturating concentrations of 200 μg/ml (n=3) or 300 μg/ml (n=1). It was observed that both antibodies were able to limit tumor growth from days 12 to 16. However, bivalent anti-PD-1 demonstrated greater efficacy in restraining tumor growth compared to monovalent anti-PD-1, as indicated by tumor volume measurements. These findings suggest that the role of mere steric hindrance in PD-1 binding to PD-L1 differs from the combined effects of steric hindrance and receptor internalization in the efficacy of checkpoint blockade for controlling tumor growth. In an anti-PD-1 resistant model, the growth of Lewis lung carcinoma tumor was unaffected by both bivalent or monovalent anti-PD-1 treatment (Fig. S3).

To elucidate the observed distinctions in the B16 F10 model, we proceeded with an analysis of the tumor-infiltrating lymphocytes (TILs). Our findings revealed that treatment with the bivalent antibody effectively increased the presence of CD45+ T-cells within the TIL population, unlike the univalent antibody (as illustrated in Fig. 5D). Additionally, we observed a similar augmentation in the abundance of CD8+ T-cells among the CD45+ TILs following bivalent antibody treatment (as shown in Fig. 5E). By contrast, the proportion of CD4+ TILs was relatively reduced in tumors treated with the bivalent antibody compared to those treated with the univalent antibody (as depicted in Fig. 5F). Taken together, these results indicate that PD-1 crosslinking and endocytosis lead to greater infiltration of CD8+ T-cells compared to CD4+ T-cells within tumors.

### Anti-PD1 steric blockade suffices to increase GZMB expression in CD8+ TILs

Our subsequent focus revolved around investigating whether different expression events were regulated by steric hindrance or internalization. Notably, we observed that both mono- and bi-valent anti-PD-1 treatments led to an increase in the expression of GZMB, a crucial cytolytic mediator, among CD8+ TILs. This increase was evident in the percentages (Fig. 5G) and mean fluorescence intensities (MFIs) (Fig. 5H). While bivalent antibody exhibited greater efficacy than univalent antibody in some experiments (2/5), this trend was not consistently observed. Instead, both mono- and bivalent anti-PD-1 consistently augmented the numbers and levels of expression of GZMB. We also analyzed several other markers, including CD44, CD62L, Tbet, TCF1, Tox, Eomes, and Ki67 (Fig. 5N). Our findings indicated some variation, with occasional increases in CD44, Tox, and Ki67 induced by both antibodies. However, these events were enhanced by both reagents and exhibited variability across experiments. Likewise, the presence and distribution of different subsets of immune cells displayed variability (Fig. 5O). Generally, the bivalent antibody induced an increase in T-cells and occasionally led to a decrease in the number of B-cells. Regarding antigens on other TILs, we tended to observe an increase in B-cells and DCs expressing MHC class II and PD-L1 (Figs. 5P and M, respectively). DCs and monocytes, on the other hand, tended to exhibit a decrease in CD80 expression (Figs. 5Q and R, respectively). However, these findings also demonstrated variability across experiments.

### Anti-PD1-induced endocytosis is needed for increased cell division and perforin expression

In contrast, among the various markers examined within the CD8+ subset, it was observed that the expression of the key cytolytic mediator perforin exhibited an exclusive increase in response to bivalent antibody rather than monovalent antibody (Fig. 5I-J). This finding was supported by standard FACs profiling (Fig. 5I) and was evident in both the percentage of CD8+ TILs expressing perforin (i.e., from 20% to 65%) (Fig. 5J) and the mean intensity of fluorescence (MIF) for perforin expression (i.e., from 1010 to 3030) (Fig. 5K). viSNE profiles further confirmed this observation, particularly within the CD8+ subset (Fig. L). Within these profiles, three distinct clusters of cells based on perforin expression were identified: cluster i with low perforin expression, cluster ii with intermediate expression, and cluster iii with high expression in untreated cells. Bivalent J43 treatment was found to enhance the expression and presence of perforin specifically in cluster i. On the other hand, monovalent anti-PD-1 did not consistently increase perforin expression in terms of the number of perforin-expressing cells or the intensity of expression (MIF values, lower panel).

Additionally, a correlation was observed between the levels of perforin expression and CD8 expression (Fig. 5M). While this correlation was not apparent in the control sample, it was observed in tumors from both monovalent and bivalent-treated mice. These findings suggest that bivalent antibody, which induces steric blockade and internalization, exerts a preferential regulatory effect on the expression of granzyme B in CD8+ T-cells compared to monovalent antibody. In other words, the combined impact of steric blockade and internalization selectively influences the cytotoxic capabilities of CD8 TILs within tumors, thereby accounting for its more potent ability to restrain tumor growth.

To explore this further, we investigated whether bivalent anti-PD1, in contrast to monovalent anti-PD1, could induce the expression of perforin in human cells within a short timeframe of 24 hours (Fig. 5S). In this experiment, human T-cells were activated for 24 hours, followed by a period of rest and incubation with either monovalent or bivalent anti-PD-1. Remarkably, bivalent Nivolumab treatment increased perforin expression, whereas univalent Nivolumab did not produce a similar effect (middle and lower panels). viSNE profiles revealed the presence of two distinct clusters of untreated CD8+ T-cells. Cluster i exhibited high levels of PD-1 expression in the untreated control CD8+ cells (middle panel), while cluster iii showed no PD-1 expression but displayed perforin expression. Treatment with bivalent Nivolumab led to the disappearance of cluster i and the emergence of a new cluster ii with low to intermediate PD-1 expression. Correspondingly, bivalent antibody induced an augmentation in perforin expression in clusters iii and ii. On the other hand, monovalent Nivolumab had no impact on promoting perforin expression and instead exhibited a pattern similar to the untreated control. Measurement analysis indicated that the mean fluorescent intensity (MFI) of perforin expression increased in response to bivalent anti-PD-1 but not univalent anti-PD-1 (lower histogram). These findings demonstrate that the bivalent dimerization of PD-1 on human cells also triggers the in vitro expression of perforin in CD8+ T-cells within a relatively short period of 24 hours. This timeframe could hold significant relevance for therapeutic interventions, as it suggests that the administration of bivalent anti-PD-1 antibodies could potentially rapidly enhance the cytotoxic activity of CD8+ T-cells.

## Discussion

Our study has explored the impact of PD-1 internalization on the effectiveness of PD-1 checkpoint blockade (ICB) in immunotherapy. We observed that conventional anti-PD-1 treatment downregulates PD-1 expression by targeting a subpopulation of CD4 and CD8 T-cells with high initial PD-1 surface receptor density, resulting in the generation of a new population of T-cells with intermediate-low expression. Further, CD8+ cytolytic T-cells were most affected, exhibiting the lowest residual PD-1 levels following down-regulation. In comparing two anti-PD-1 drugs, Nivolumab displayed superior proficiency in downregulating PD-1 expression in human cells compared to Pembrolizumab. Most importantly, we discovered that anti-PD-1 internalization was dependent on PD-1 receptor crosslinking. While both monovalent antibodies (which facilitated steric blockade) and bivalent antibodies (which promoted steric blockade and endocytosis) limited tumor growth, the bivalent antibody routinely surpassed the monovalent antibody in reducing tumor size. At the molecular level, while both mono and bivalent antibodies increased the expression of granzyme B, only bivalent antibody increased the expression of the second key cytolytic pore protein, perforin. These findings unveil a novel mechanism that extends beyond steric blockade in checkpoint blockade, emphasizing the potential of targeting PD-1 endocytosis to optimize checkpoint blockade for anti-tumor immune responses.

Our initial observations revealed that all antibodies used in checkpoint blockade, whether in mouse (J43, RPMI) or in human (Nivolumab and Pembrolizumab), induce the internalisation of PD-1. Unexpectedly, the effects were observed primarily on T-cells expressing high levels of PD-1 leading to the generation of a population of cells with low to intermediate PD-1 expression. This was particulary evident in viSNE analysis where we observed the visible loss of clusters with high PD-1 expression accompanied by emergence of a new clusters of CD4 and CD8 T-cells with intermediate PD-1 expression. On a broad scale, approximately 50-60% of PD-1 molecules were internalized from the cell surface. The reason for this selectivity is not entirely clear, it is likely related to the need for crosslinking of PD-1 for internalization. Once the receptor density falls below a certain threshold, the density of receptors is insufficient to support effective crosslinking between adjacent PD-1 receptors for internalization. Alternatively, it is possible that certain receptors are resistance to internalization, aligning with the concept of spare receptors reaching maximum response before receptor saturation. Overall, these findings demonstrate that anti-PD-1 checkpoint blockade involves more than simple steric interference of PD-1 binding to PD-L1, as it operates in conjunction with the active removal of PD-1 from the surface of T-cells.

In terms of anut-human antibodies, nivolumab was routinely found to bind at higher levels of activated human T-cells and to be more effective in down-regulating PD-1 from the cell surface than pembrolizumab. This was seen in CD4 and CD8+ T-cells and in terms of the rate and level of internalisation. This may be related to the binding of each antibody to different sites on the PD-1 receptor. An unexpected N-terminal loop in PD-1 dominates binding by Nivolumab ^77^. Pembrolizumab binding may involve a unique and large patch of interactions engaging the C’D loop of PD-1 ^68^. Whether this difference in the efficacy of internalisation affects the killing of cancer targets awaits future experiments. In ether case, we found that anti-PD-1 internalisation affected CD4 and CD8+ T-cells as well as their effector memory and effector subsets. In this sense, there was no inherent predisposition for any specific subset, such as memory or effector subset, to downregulate the receptor. However, at the same time, surprisingly, we observed less binding to effector CD8+ T-cells relative to effector memory cells. Further, the downregulation on CD8, and particularly CD8 effectors, resulted in a smaller residual level of PD-1 expression than seen on CD4+ T-cells. This observation aligns with the finding that anti-PD-1 ICB primarily affects CD8 effector T-cells ^11^. In this model, the preferential effect could in part be related to the greater reduction of residual PD-1 on the surface of CD8+ cells.

Mechanistically, our findings revealed that PD-1 downregulation required receptor crosslinking. Monovalent anti-PD-1 antibodies, such as J43 for mice and Nivolumab for humans, were unable to downregulate the receptor. Instead, the same bivalent antibody was effective. Importantly, this characteristic enabled us to investigate the impact of steric hindrance mediated by monovalent antibodies relative to the combined effects of steric hindrance and receptor internalization as mediated by bivalent antibdies in tumor immunotherapy. Both antibodies exhibited comparable binding affinity to T-cells, as demonstrated through a titration ranging from 0.01 to 5 μg/ml during incubation. High levels of antibody (200-300 μg/mouse) were administered every 2 days over a span of 20 days. Under these conditions, both monovalent and bivalent antibodies impeded tumor growth, emphasizing the significance of steric blockade. However, the bivalent antibody exhibited greater effectiveness (n=4), each experiment consisting of five mice per treatment group. In most cases, this efficacy of tumor regression was correlated with a relative increase in the proportion of CD8+ versus CD4+ TILs. Consistent variations in other cell populations, such as dendritic cells (DCs) or polymorphonuclear leukocytes (PMNs), were not observed. In addition, consistent differences in other markers, such as CD44 and CD62L, were not detected. In the case of a PD-1 resistant tumor such as LLC, netiher mono-valent nor bivalent antibody was offective.

One parameter that was consistently shared on CD8+ T-cells in response to monovalent and bivalent antibodies was the cytolytic mediator GZMB on CD8+ T-cells. This is a prime effector required for the killing of tumor cells and could account for the protective effects against tumor growth are needed to accommodate the increased expression of GZMB ^78^. The fundamental property therefore appears linked to the steric blockade effect of anti-PD-1. By contrast, the action of bivalent antibody could be distinguished from the monovalent antibody with the former consistently upregulating perforin expression. Perforin is a critical protein involved in the immune response, particularly in the cytotoxic activity of CD8+ T-cells ^78^. It plays a crucial role in the elimination of virally infected or cancer cells, by creating pores in their cell membranes and then facilitating the entry of cytotoxic molecules. This consistent pattern was observed in TILs across all experiments. Interestingly, the effects could both *in vivo* and in *in vitro* activation using preactivated PD-1 expressing human cells that were subsequently rested, and incubated with the antibody for only 24-48 hours. Further, it was observed for bivalent J43 in mouse and Nivolumab in human T-cells. This observation highlights the importance of receptor mediated crosslinking and removal of PD-1 from the cell surface in the induction of perforin expression. of the structural conformation of PD-1 in modulating its functional outcomes. It might be linked to a threshold effect where the steric blockade combined with PD-1 removal from the cell surface reduces negative signaling more effectively that impair T-cell activation. Another possibility is that receptor crosslinking could induce downstream signaling pathways involved in perforin expression compared to the monovalent interaction. In this later context, receptor crosslinking is known to be essential for signaling through multiple receptors ^79,80^. Other possible mechanisms could involve internalized endosomes that can transmit signals, as exemplified by the continued activation of the internalized tyrosine kinase receptor TrkA in signaling endosomes ^81^. The precise contribution of each mechanism, or whether both mechanisms work in tandem to generate signals, remains to be elucidated. Either way, our findings unveil a novel mechanism in checkpoint blockade where steric blockade combined with the removal of PD-1 from the cell surface by endocytosis can complement and optimize therapy via a perforin mediated mechanism. In this way, the targeting of PD-1 internalisation holds promise for enhancing anti-tumor immunity and improving PD-1 checkpoint blockade therapy.

Lastly, our study provided information on how PD-1 is internalised into cells. PD-1 endocytosis was seen to be eventually incorporated into early and late endosomes as shown by EEA1 (Rab5a) and Rab7a colocalization. Further small amounts could be re-cycled to the cell surface. Although sucrose was found to inhibit internalization, neither inhibitor, Dynasore nor Pitstop2 affected PD-1 internalisation, using human or mouse cells and different assays. This despite observable effects on CD28 or MHC class 1 internlisation. These observations suggest that conventional clathrin or caveolae-mediated endocytosis pathways are not major mediators of PD-1 internalisation. In this context, a new mobile endocytic network connecting clathrin-independent receptor endocytosis to recycling has been reported which promotes T cell activation ^75^. Further we observed no detectable ubiquination despite seeing the presence of multiple ubiquinated bands in cell lysates from cells. Our antibody induced effects therefore clearly operate deifferently from the E3 ligase FBXO38 ubiquitination mechanism that has been reported to control steady-state PD-1 turnover ^64^. Another possible pathway involves flotillin-1 as a determinant of a clathrin-independent endocytic pathway in mammalian cells ^82^ or possibly a mechanism involving the AP2 complex as we previously reported for CD28 and CTLA-4 ^59,83^. However, the receptor nevertheless was found in an internalized form to co-localize with the early endosome marker EEA1 (Rab5a) and the late endosome marker Rab7a and is therefore eventually incorporated into the endosomal pathway.

Future studies will be necessary to further elucidate the relative benefits of enhanced PD-1 signaling in different clinical scenarios. Some studies have indicated that Nivolumab leads to better overall survival outcomes in this specific patient subgroup compared to Pembrolizumab. Additionally, Nivolumab has shown effectiveness in advanced renal cell carcinoma, including patients who have previously received anti-angiogenic therapy, with better overall survival benefits observed in certain studies when compared to alternative treatments, including Pembrolizumab. While Pembrolizumab has also demonstrated activity in Hodgkin lymphoma, Nivolumab has a specific approved indication in this context. Overall, these findings contribute to our understanding of the intricate interplay between PD-1 signaling, perforin expression, and CD8+ T-cell function. They provide valuable insights into the potential mechanisms by which bivalent anti-PD-1 antibodies can influence immune responses and pave the way for further investigations and therapeutic advancements in the field of immunotherapy.

## Materials and Methods

### Cells and Reagents

Blood samples of healthy donors were provided by Héma-Québec as LRS (LeucoReduction System) chamber. Peripheral blood mononuclear cells (PBMC) were isolated from human samples using Ficoll-Paque density centrifugation (Corning, Fisher Scientific, CA). PBMC were cultured in RPMI 1640 medium (Corning, Fisher Scientific, CA) supplemented with 10% heat-inactivated FBS (Fetal Bovine Serum), 1% glutamine (200mM) and 1% penicillin/streptomycin (10,000U/ml) (Gibco, ThermoFisher, CA) (complete RPMI media). Cells were maintained at 37°C in an atmosphere of 5% CO_2_. To express PD-1 receptor, cells were stimulated with soluble anti-human CD3 (clone OKT3) and anti-human CD28 (clone 9.2) at 3 and 2µg/ml respectively, for 2 days. Concanavalin A (Con A) (Sigma Aldrich, CA) was also used at 10 µg/ml for 2 days to activate cells, which were then washed with complete RPMI1640 medium and maintained overnight in 5% CO_2_ incubator.

### Antibodies and chemicals

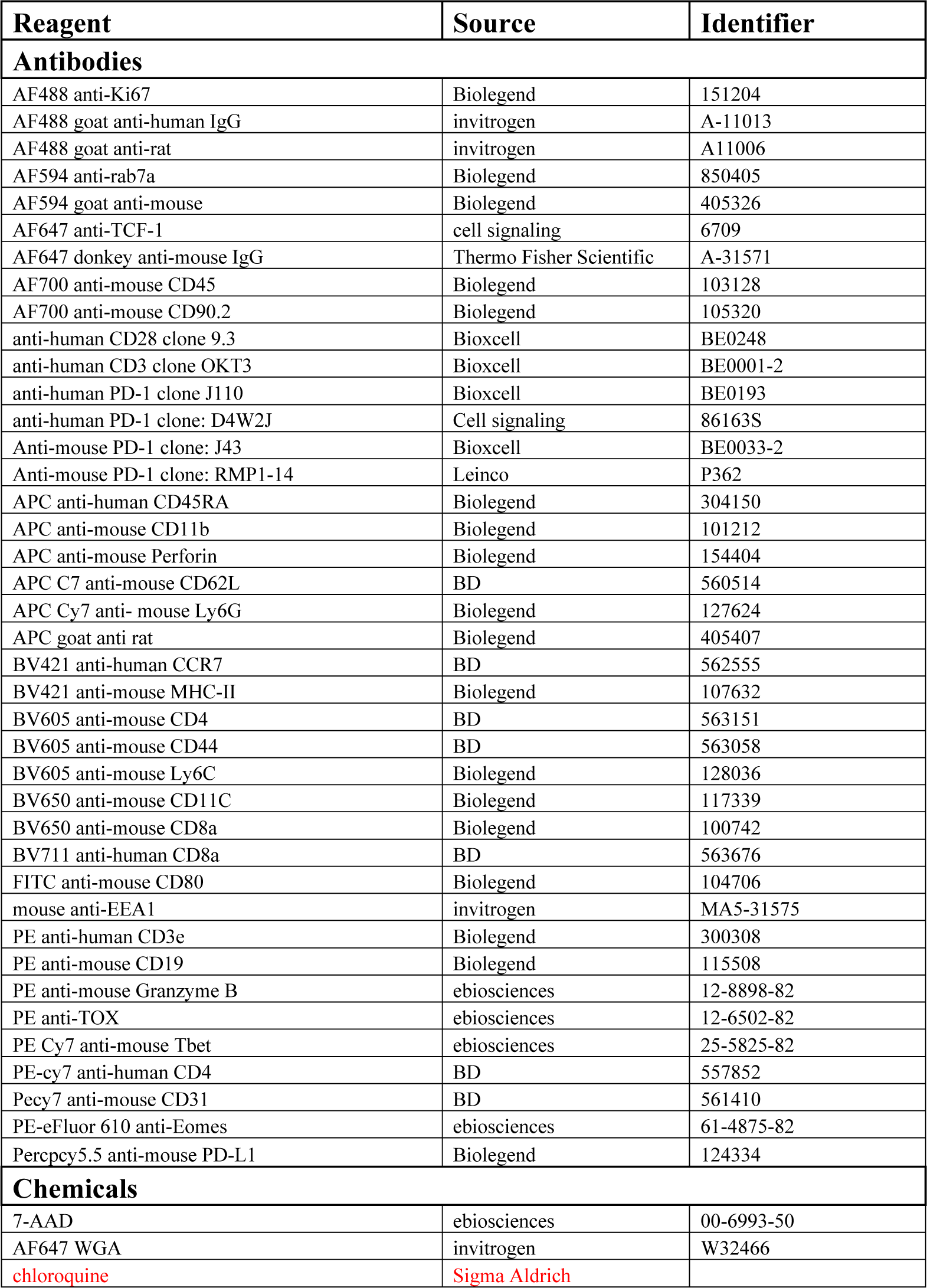
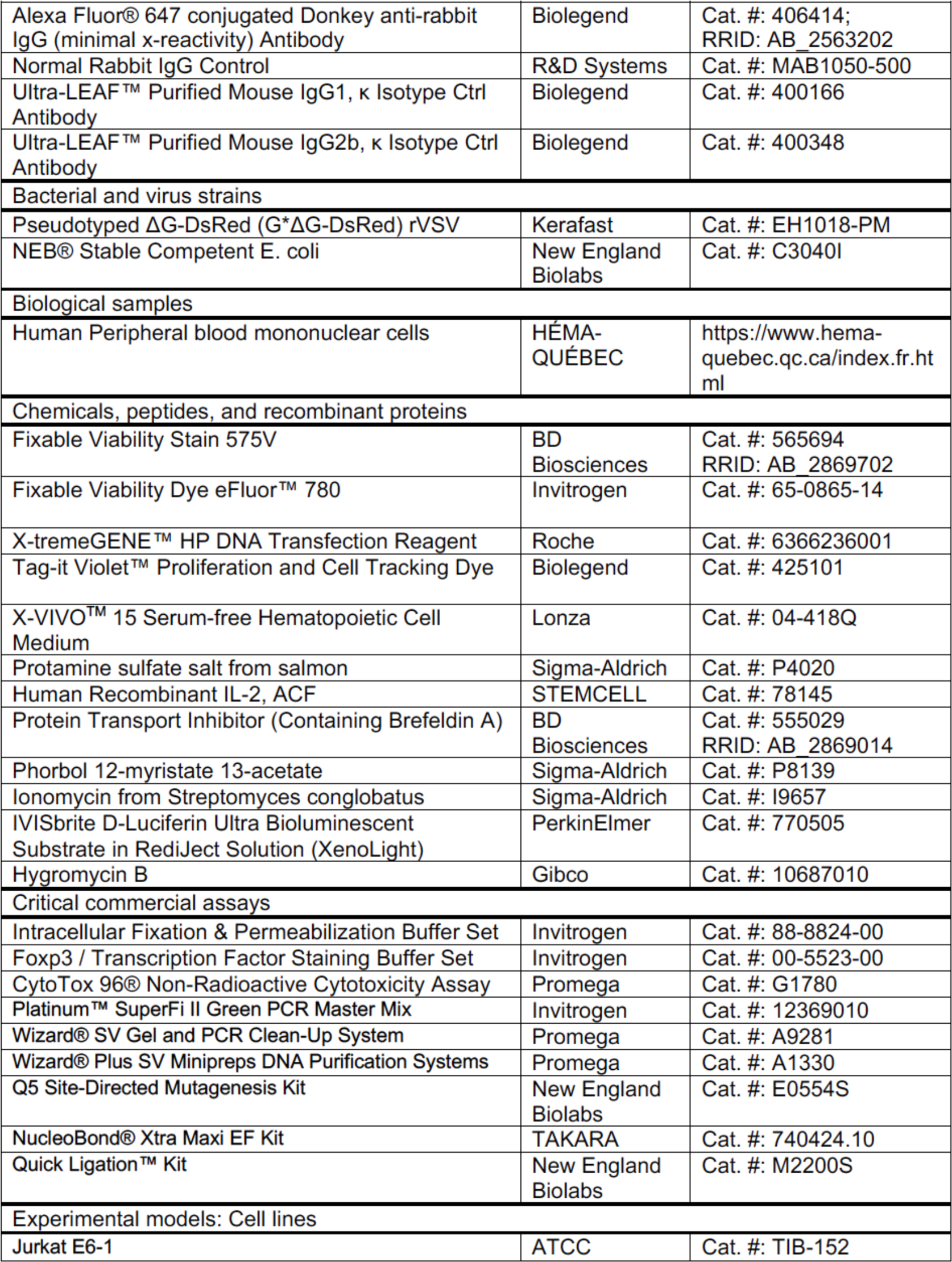
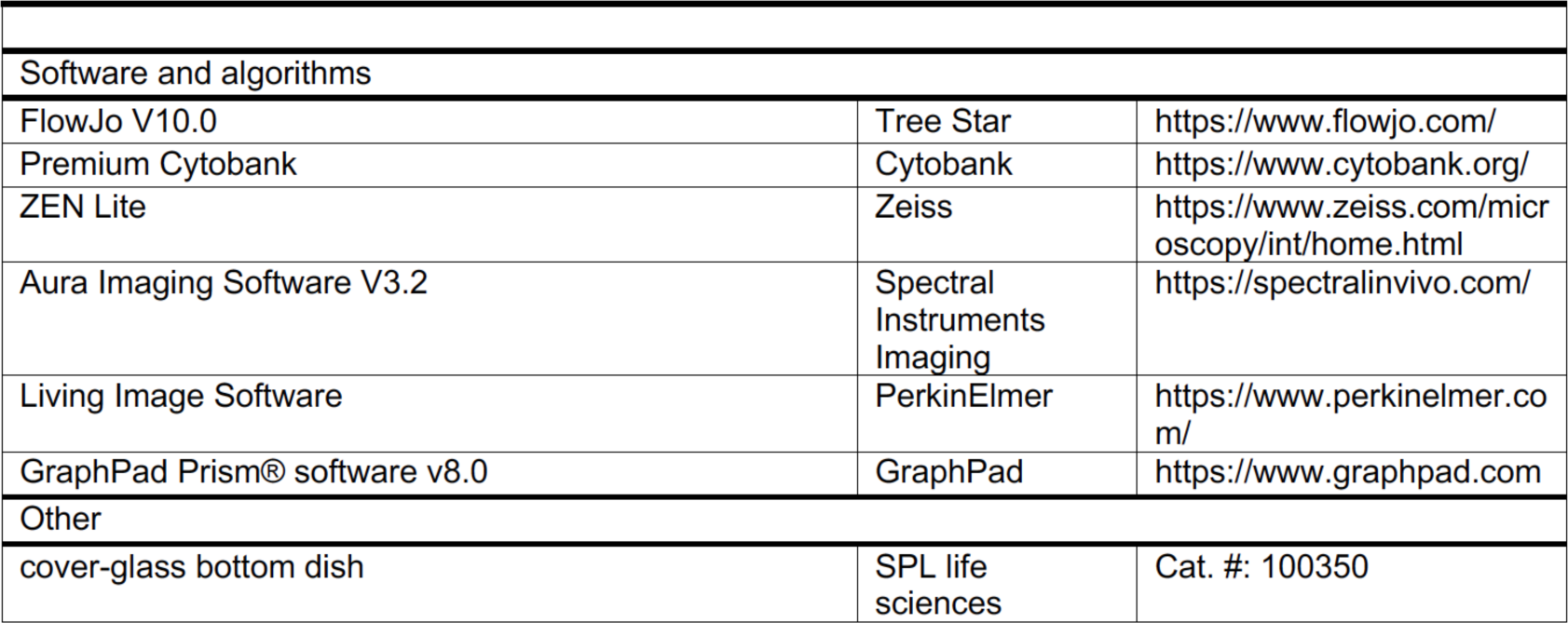

### Internalization and down-modulation assays

To assess the internalization of PD-1 and CD28 in human T cells, we followed the methodologies outlined in Céfai et al. (1998) and Walter et al. (2008). Human peripheral blood mononuclear cells (PBMCs) were activated using either anti-CD3/CD28 or Con A. To block Fc receptors, the cells were treated with PBS containing 10% donkey serum for 20 minutes at room temperature. Subsequently, the cells were washed and incubated with 1µg/ml of anti-human PD-1 monoclonal antibody (mAb) for 30 minutes at 4°C. After three washes with cold PBS, we added AF647-conjugated donkey anti-mouse IgG antibody in PBS containing 1% donkey serum. Following a 30-minute incubation on ice, the cells were washed and divided into multiple tubes (typically 2.5-3×106 cells per tube) and incubated at 37°C for various durations. One set of cells was kept on ice to represent the total cell-associated fluorescence, while another set was incubated with acidified PBS (pH 2.0) supplemented with 0.03M sucrose and 10% FBS to indicate intracellular fluorescence. The cells were then washed three times with cold RPMI 1640 containing 10% FBS and once with FACS buffer. Additionally, one sample was incubated solely with AF647-conjugated anti-mouse IgG antibody as a negative control.

For PBMCs activated with Con A, the cells were also stained with PE-anti-human CD3 antibody for 30 minutes on ice. To exclude nonviable cells, all samples were stained with 7-AAD and analyzed using flow cytometry. For CD28 internalization, freshly isolated PBMC were incubated with 1 µg/ml anti-human CD28 Ab for 30min on ice. After washing with cold PBS, AF647 conjugated anti-mouse IgG Ab was added. The next steps were the same as described in PD-1 internalization.

For the down-modulation assays, following the Fc receptor blocking step described earlier, human peripheral blood mononuclear cells (PBMCs) activated with Con A were exposed to 1µg/ml of either anti-human PD-1 (specifically J110) or anti-human CD28 (9.2) monoclonal antibodies (mAbs) for 30 minutes on ice. The cells were then washed and suspended in cold RPMI 1640 medium. They were divided into multiple tubes, with approximately 2.5×106 cells per tube, and incubated at 37°C for various durations. At the designated time points, the samples were washed with cold PBS and incubated with 1µg/ml of AF647-conjugated anti-mouse IgG antibody for 30 minutes on ice. This allowed the detection of any remaining PD-1 or anti-PD-1 on the cell surface. After washing with cold FACS buffer, the cells were stained with PE-anti-human CD3 for 30 minutes on ice. Subsequently, the cells were washed, stained with 7-AAD, and analyzed using flow cytometry.

For the downmodulation involving the use of inhibitors of endocytosis, murine T-cells were activated for 48h with Con A 2.5 ug/ml, washed, and rested overnight. Cells were treated with either DMSO, Pitstop2 (15 uM, clathrin-dependent endocytosis), or Filipin III (2.5 ug/ml, inhibitor of caveolae-mediated endocytosis) for 1h at 37°C in complete media. Cells were then washed and incubated with anti-PD-1 (clone: RMP1-14; 20 ug/ml) on ice for 1h in complete media. Cells were washed and then kept on ice or incubated at 37°C for 1h in complete media. The cells were stained with secondary APC-conjugated anti-rat, then washed and stained with AF700 anti-CD90.2, BUV395 anti-TCRb, BV605 anti-CD4 and BV650 anti-CD8a.

For humanized nivolumab and pembrolizumab treatments, cells were exposed to a concentration of 1µg/ml of the respective antibodies for 30 minutes on ice. As a negative control, cells were solely incubated with AF488-conjugated anti-human IgG secondary antibody. At the designated time points, samples were collected on ice, washed with cold PBS, and then incubated with 1µg/ml of AF488-conjugated goat anti-human IgG antibody (previously diluted in PBS containing 1% goat serum) for 30 minutes on ice. Subsequently, the samples were washed twice with cold PBS and stained with fixable viability stain 510 (FVS 510) for 20 minutes on ice. The cells were then washed and stained for 30 minutes with the indicated antibodies: PE-anti-CD3, PE-Cy7-anti-CD4, BV711-anti-CD8, APC-anti-CD45RA, and BV421-anti-CCR7. Following another round of washing with FACS buffer, the samples were fixed in 2% paraformaldehyde (PFA) before undergoing flow cytometry analysis. Fluorescence minus one (FMO) controls were included to establish positive thresholds.

### Generation of Monovalent Antibodies

Monovalent anti-PD-1 were generated from the bivalent anti-PD-1 mAbs (clone-RMP1-14) using the PierceTM Fab and Fabc Micro Preparation Kit according to the manufacturer’s instructions. Briefly, the disulfide bonds connecting the two heavy chains of RMP1-14 were reduced using 2-mercaptoethylamine-HCl (catalog number 20408) at 37°C for 1 hour. Approximately 3mg of RMP1-14 was used per bottle of 2-Mercaptoethylamine-HCl (6 mg), a mild reducing agent commonly used to selectively cleave antibody hinge region disulfides and separate the antigen binding fragments (Fabs and Fabcs). Following the antibody reduction step, the solution was then passed through PBS-EDTA-pre-charged ZebaTM Spin Desalting Columns (Thermo Fisher Scientific, catalog number 89889) where samples were buffer exchanged into the cleavage buffer provided in the kit. The Zeba columns contain a proprietary size exclusion resin that enables buffer exchange and removal of salts through centrifugation of RMP1-14 antibody was added to a 2mL Zeba column and spun at 1500 x g for 2 minutes to elute the buffer exchanged sample. The monovalent anti-PD-1 Fab fragments were then analyzed by SDS-PAGE under non-reducing and reducing conditions to confirm cleavage of the antibody hinge region and separation of heavy and light chains from the parent bivalent RMP1-14 antibody.

### Tumor growth assays

C57BL/6 mice were housed at the Hôpital Maisonneuve Rosemont animal facility (Montreal, QC, Canada). Mice were housed in individually ventilated cages (IVC) and all experiments were approved by the CR-HMR Ethical Approval (Le Comité de protection des animaux du CIUSSS de l’Est-de-l’Île-de-Montréal (CPA-CEMTL), F06 CPA-21061 du projet 2017–1346, 2017-JA-001/2). Mice, aged 7–8 weeks, were implanted intradermally with 50,000 B16-F10, B16-PD-L1 melanoma cells that overexpress PD-L1 or LCC tumors. B16 F10 melanoma tumor cells and LLC tumor cells were obtained from American Type Culture Collection (ATCC). B16-PD-L1 melanoma cells were kindly provided by Dr. M. Ardolino, Ottawa, Canada. The cells were cultured in Dulbecco’s modified Eagle’s medium (DMEM; Gibco) supplemented with 10% fetal bovine serum (FBS; Gibco) and 1% penicillin/streptomycin (P/S; Gibco). Cells were incubated at 37°C in a humidified atmosphere containing 5% CO2. When cells reached approximately 70% confluency during the logarithmic growth phase of in vitro cultivation, they were prepared for injection. The culture medium was aspirated and the cells were washed twice with 1X phosphate-buffered saline (PBS; Gibco) to remove residual serum. Trypsin-EDTA (Gibco) was used to detach the cells from the culture flask surface. FBS was added to neutralize the trypsin and the cells were collected by centrifugation at 200 x g for 5 minutes. The cell pellets were resuspended in PBS and cell counts were determined using a hemacytometer. Only single-cell suspensions with >90% viability by trypan blue exclusion were used for injection. B16-F10 is a mouse melanoma line while Lewis lung carcinoma cells are hypermutated Kras/Nras–mutant^84^

### Isolation of tumor infiltrating lymphocytes (TILs)

Solid tumors were harvested from mice 16-20 days post tumor challenge under sterile conditions. Tumors were collected and measured to ensure they reached approximately 100-200 mm3 in size. Tumors were placed in sterile phosphate-buffered saline (PBS) on ice. Tumors were manually disrupted and minced into small pieces (<1 mm3) using sterile surgical blades and dissection scissors. Minced tissue was then digested in RPMI 1640 medium supplemented with 5% fetal bovine serum (FBS), 1% Penicillin-Streptomycin, 200 U/mL collagenase type IV (Gibco), and 30 U/mL DNase I (Sigma) at 37°C for 1 hour with periodic agitation. Following digestion, the cell suspension was passed through a 70 μm cell strainer into a 50 mL conical tube to obtain a single-cell suspension and remove remaining tumor fragments and debris. The cell suspension was then centrifuged at 300xg for 8 minutes at 4°C. The cell pellet was resuspended in 40% ficoll-paque (GE Healthcare) and overlaid carefully on top of 60% ficoll-paque solution. Tubes were centrifuged at 700xg for 20 minutes at room temperature with the brakes off. The lymphocyte layer containing tumor infiltrating lymphocytes (TILs) was collected from the interface between the ficoll layers and transferred to a new 50 mL conical tube. TILs were washed twice with cold RPMI medium by centrifuging at 300xg for 8 minutes at 4°C to remove residual ficoll. TILs were then counted, assessed for viability by trypan blue exclusion, and either used immediately for staining or cryopreserved in liquid nitrogen until further use.

### Flow cytometry

Flow cytometry analysis involving antibody staining of surface receptors was performed using a suspension of 106 cells in 100μl of PBS. The cells were then exposed to a 1:100 dilution of the primary antibody at 4oC for a duration of 2 hours. Subsequently, the cells were washed twice with PBS. In certain cases, the cells were suspended in 100μl of PBS containing a secondary antibody and incubated for an additional 1 hour at 4oC. Cell staining data was analyzed using a Beckman Coulter CytoFLEX S flow cytometer and the CytExpert software. For intracellular staining, cells were fixed using 4% paraformaldehyde (PFA) and permeabilized with 0.3% saponin (Sigma-Aldrich). The cells were then stained with the desired antibody in PBS containing saponin for 2 hours at 4oC. If primary antibodies were not conjugated, a secondary antibody incubation was carried out subsequently.

### Visualisation of internalised receptors by fluorescence microscopy

PD-1 internalization was also conducted by fluorescence microscopy. Briefly, activated cells were incubated with 1µg/ml of anti-PD1 Ab (J110) for 30 minutes on ice, followed by a washing step and incubation with AF647-anti-IgG secondary Ab. Cells were then washed, resuspended in cold RPMI 1640 and incubated at 37°C for 30min or at 4°C for 90min. After incubation, cells were washed in cold PBS and divided into two parts: one was kept on ice while the other was acid-stripped with PBS (pH2) for 5 min and washed 3 times in cold RPMI 1640/10%FBS. Samples (45×10^4^/tube) were then washed with cold PBS, fixed in 2% PFA, and centrifuged by cytospin at 600rpm for 7 min. Subsequently, slides were washed for 2 min in 70% followed by 100% ethanol, and 10µl of Vectashield anti-fade mounting medium (containing 1µg/ml of DAPI) (Vector laboratories) was added on the cells spot. Then, coverslips were sealed by nail polish before visualization in fluorescence microscopy.

Further, for an assessment of anti-PD1 antibody complex co-localization with the early endosome marker EEA1 (Rab5a) and the late endosome marker Rab7a, pre-activated 48hr C57BL6J mice-derived splenic T-cells were incubated with anti-PD-1 (clone RMP1-14) for 1h followed by treatment with acidic PBS (pH: 2) to remove cell surface antibody followed by fixation and the staining with either mouse anti-EEA1 followed by AF594-anti mouse or AF594-Rab7a Abs.

### Recycling determination

Activated PBMCs expressing PD-1 were incubated with primary Ab for 30min on ice, followed by washing in cold PBS and incubation with AF488-anti human IgG or AF647-anti-mouse IgG secondary Abs, respectively. After 30 minutes on ice, samples were washed twice in cold PBS, resuspended in cold complete RPMI 1640 medium, and incubated for 60 minutes at 37°C. One sample was left on ice for 60 minutes to determine the starting level of PD-1 expression and to control the efficacity of acid stripping. As a negative control, cells were incubated only with secondary Abs.

Subsequently, samples were washed twice with cold complete RPMI1640, acid stripped with PBS (pH:2) and resuspended in warm complete medium in the absence or presence of 100 µM chloroquine (CQ) (Sigma Aldrish, CA). Cells were then transferred to a 12-well plate (4×10^6^/well) and re-incubated at 37°C in 5% CO_2_ for the indicated time periods (0, 12h, 60h). After each time point, cells were collected, washed twice in cold PBS, left untreated, or acid-stripped where indicated (in the presence of CQ and MG132). Next, all samples were stained with 7-AAD before flow cytometry analysis.

### Drug inhibition of endocytosis

To evaluate the effect of dynamin inhibitor on PD-1 and CD28 internalisation, activated PBMCs with Concanavalin A were incubated with 1 µg/ml of Nivolumab or anti-hCD28 Ab for 30 minutes on ice. Cells were then washed and incubated with 1 µg/ml AF488-anti human IgG or AF647-anti-mouse IgG Abs, respectively, for 30 minutes on ice. Next, samples were washed twice in cold PBS, resuspended in serum-free RPMI medium in the presence of DMSO or Dynasore (100 µM) (ab120192, Abcam), and incubated at 37°C from 0 to 90 min. After each time, cells were washed and stripped or not with acidified PBS (pH 2). All samples were then stained with PE-anti-CD3 for 30 min on ice, washed, and analyzed by flow cytometry after the addition of 7-AAD.

To evaluate the effect of Pitstop2 on PD-1 downmodulation, activated splenocytes with Con A were incubated with 15 µM Pitstop2 or DMSO for 15 min at 37°C. After the cells were washed, they were incubated with 1 µg/ml of anti-mouse PD-1 (clone RMP1-14). Next, samples were washed twice in cold PBS, resuspended in complete with 15 µM Pitstop2 or DMSO, and incubated at 37°C from 0 to 60 min. Subsequently, samples were stained with APC-conjugated anti-rat IgG, CD4, CD8 and TCRb and analyzed by FACs.

To evaluate the effect of Pitstop2 on MHCI-H2K^b^ endocytosis, activated splenocytes with Con A were incubated with 15 µM Pitstop2 or DMSO for 15 min at 37°C. After the cells were washed, they were incubated with 1 µg/ml of FITC-conjugated anti-mouse MHCI-H2K^b^ (Cat# 116505, Biolegend). Next, samples were washed twice in cold PBS, resuspended in complete with 15 µM Pitstop2 or DMSO, and incubated at 37°C for 60 min. After this incubation, cells were washed and stripped or not with acidified PBS (pH 2) and analyzed by flow cytometry.

### Immunoprecipitation and Western blotting

To carry out an immunoprecipitation, 25 x10^6^ cells were lysed in 500 µl of 1% Triton X-100 lysis buffer (1% Triton X-100, 25 mM Tris pH 8, 150 mM NaCl, 2 mM Na₃VO₄, 10 mM NaF) containing a mixture of protease inhibitor (Halt™ Protease Inhibitor Cocktail, Thermofisher) as described^85,86^. After incubation for 30 min at 4°C, lysates were pelleted and supernatants were used for immunoprecipitation and/or western blot directly. For immunoprecipitation, 2.5 µg of rabbit anti-PD-1 (clone D4W2J, 86163S Cell signaling) or rabbit IgG (ab6709, Abcam) were coupled to Protein A-Sepharose beads for 3h at 4°C. Lysates were then incubated with antibody-coupled beads and rotated at 4°C for 1 h. Beads were washed three times in cold lysis buffer, they were resuspended in 70 µl of loading buffer (2X) and boiled for 10 min at 95°C. Proteins were separated on 10% SDS-PAGE and transferred to nitrocellulose for immunoblotting. Proteins were transferred from the gel onto a nitrocellulose membrane using an electric current and fixed to the nitrocellulose and probed with antibodies against the target proteins (anti-PD and anti-ubiquitin; Invitrogen Ubi-1, Cat #13-1600). Bound antibody was revealed with horseradish peroxidase-conjugated rabbit anti-mouse antibody using enhanced chemiluminescence (Amersham Biosciences).

## Data analysis

Results from the different studies were presented as means ± standard deviation (SD) from at least three independent experiments. Flow cytometry data were analysed by FlowJo software (version 10). For viSNE plots, data were exported from FlowJo as FCS files to Cytobank premium and viSNE analysis was performed with perplexity set to 50. Graphs and statistical analysis were performed using GraphPad Prism software (version 8). The rate of PD-1 internalization at an indicated time (tx) was determined as follows: (MFI of acid-treated samples at tx / MFI of untreated samples at tx) x100. The rate of PD1 or CD28 down-modulation was determined as follows: (MFI of samples at tx/ MFI of samples at t0) x100. The study of PD-1 internalization by fluorescence microscopy was performed by Zeiss widefield imaging system using AxioVision software. Images were processed using Fiji software (Fiji is just ImageJ, version 1.0).

## Funding

C.E.R. was supported by the Canadian Institutes of Health Research Foundation grant (159912).

## Author Contributions

Conceptualisation: CER, AO, NPA, MB and EBS.

Investigation: EBS, MB, NPA and AO conducted most experiments.

Methodology: CER coordinated arrangements for the processing of samples. EBS and CER coordinated access to healthy donors from HemaQuebec (agreement between the CER and HemaQuebec). CER coordinated ethical approval to work with samples at the HMR.

Supervision: CER

Writing: C.E.R. wrote the manuscript with assistance from EBS, NPA and AO.

## Competing interests

The authors declare no competing interests.

## Data and material availability

All data, code, and materials used in the analyses will be available to any researcher for purposes of reproducing or extending the analyses.

## Supplementary Figures

**table S1:**
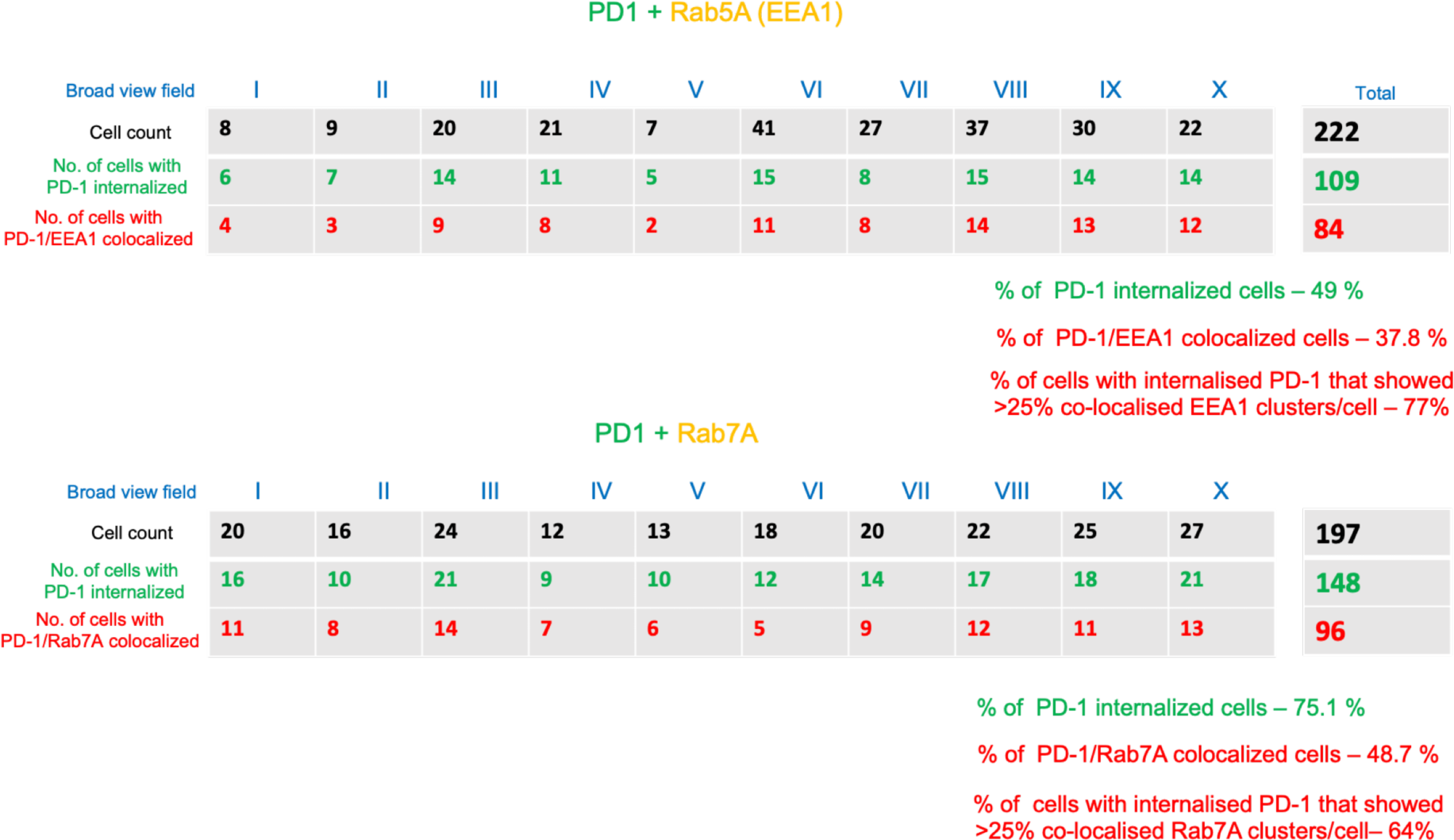
Table showing the numbers of T-cells with internalized PD-1, percent with PD1/ Rab5A (EEA1) or Rab7A colocalization and percent of cells in which greater than 25% of PD-1 clusters colocalized with EEA1 or Rab7A. Upper panel: PD1 and Rab5A (EEA1); lower panel: PD1 and Rab7A (n=3).

| Broad view field | I | II | III | IV | V | VI | VII | VIII | IX | X | Total |
| --- | --- | --- | --- | --- | --- | --- | --- | --- | --- | --- | --- |
| Cell count | 8 | 9 | 20 | 21 | 7 | 41 | 27 | 37 | 30 | 22 | 222 |
| No. of cells with PD-1 internalized | 6 | 7 | 14 | 11 | 5 | 15 | 8 | 15 | 14 | 14 | 109 |
| No. of cells with PD-1/EEA1 colocalized | 4 | 3 | 9 | 8 | 2 | 11 | 8 | 14 | 13 | 12 | 84 |

| Broad view field | I | II | III | IV | V | VI | VII | VIII | IX | X | Total |
| --- | --- | --- | --- | --- | --- | --- | --- | --- | --- | --- | --- |
| Cell count | 20 | 16 | 24 | 12 | 13 | 18 | 20 | 22 | 25 | 27 | 197 |
| No. of cells with PD-1 internalized | 16 | 10 | 21 | 9 | 10 | 12 | 14 | 17 | 18 | 21 | 148 |
| No. of cells with PD-1/Rab7A colocalized | 11 | 8 | 14 | 7 | 6 | 5 | 9 | 12 | 11 | 13 | 96 |

% of PD-1 internalized cells – 75.1 %
% of PD-1/Rab7A colocalized cells – 48.7 %
% of cells with internalised PD-1 that showed >25% co-localised Rab7A clusters/cell– 64%

**Figure S1:**
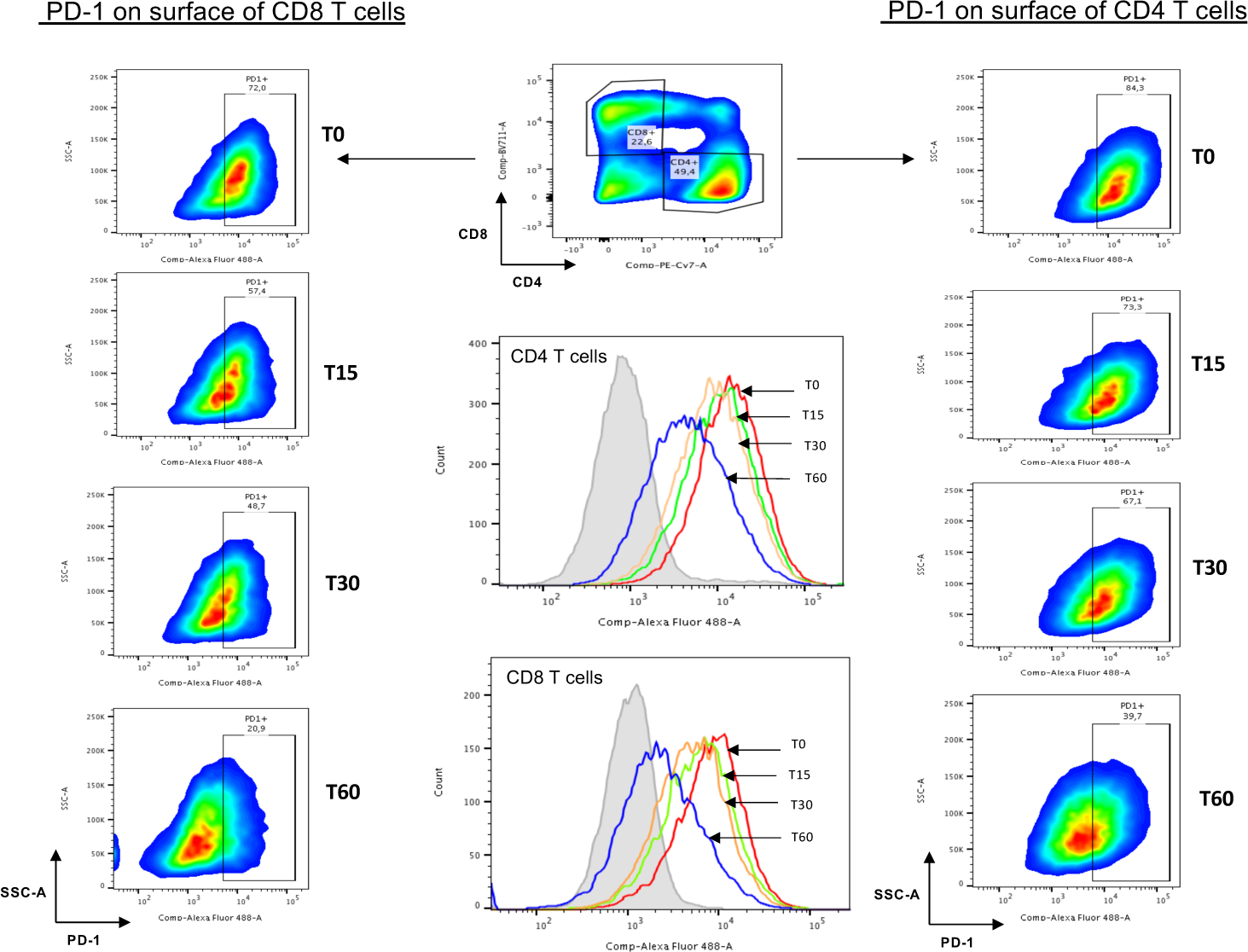
Cytometry analysis of PD-1 axpression on CD4 and CD8+ T-cells. The figure illustrates the gating and images of the impact of nivolumab on the down-regulation of PD-1 expression in effector memory CD8 (left panel) and CD4+ (right panel) T-cells. The middle panel shows the histograms showing the reduction in PD-1 expression. The representative experiment was conducted 4 times (n=4).

**Figure S2:**
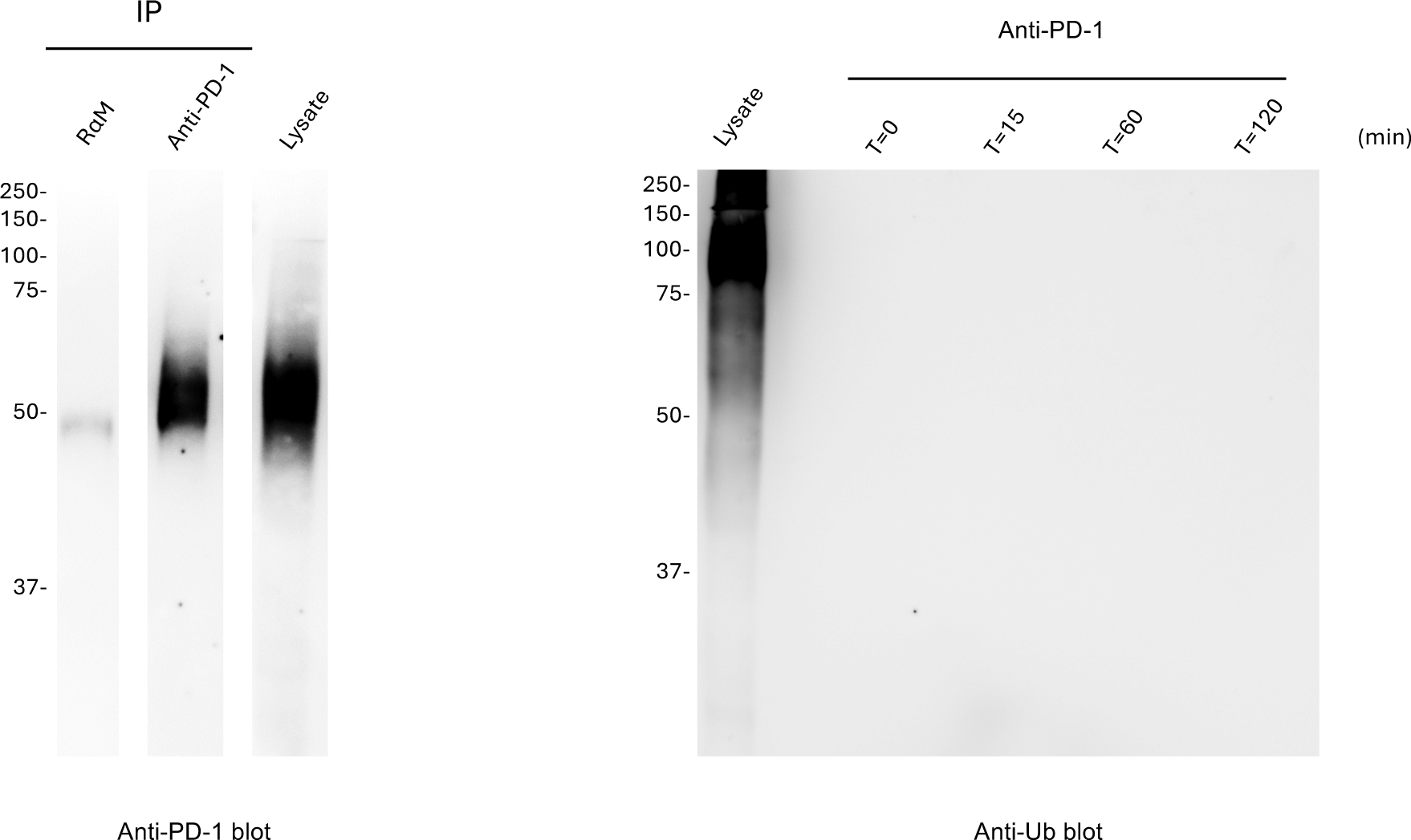
Full length blots for anti-PD1 and anti-Ub blotting (match for Figure 4H).

**Figure S3:**
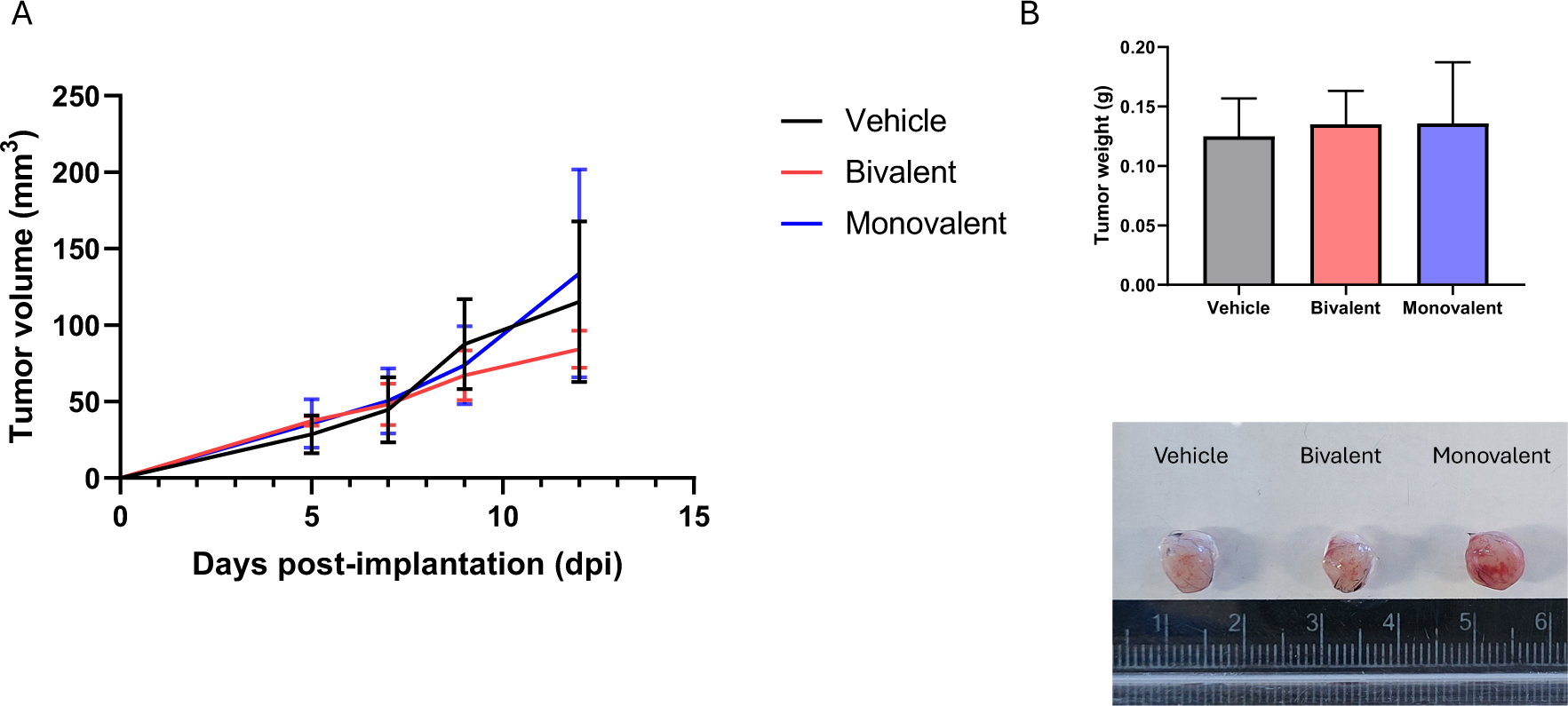
Histogram showing the growth of Lewis lung carcinoma (LLC) tumors in mice in the absence and presence of biovalent and monovalent anti-PD-1. LLC cells were expanded in vitro following inplantation in C57/Bl mice subcutaneously followed by growth of tumor until day 6 at which time mice were injected every 2^nd^ day with either bilvalent (red) or monovalent (blue) anti-PD-1 (3 mice/group). Panel A: Tumor growth curve for the LLC tumor. Neither bivalent nor monovalent anti-PD-1 altered tumor growth. Panel B: Upper panel: Tumor weight of LLC tumors on day 12. Lower panel: Example of images of tumors treated with vehicle (PBS), monovalent or bivalent antibody.

## Notes

### Competing Interest Statement

The authors have declared no competing interest.

